# Sensory experience drives glutamate spillover, presynaptic LTP and functional connectome remodeling in the barrel cortex

**DOI:** 10.64898/2026.08.22.746422

**Authors:** Olga Kopach, James P. Reynolds, Thomas P. Jensen, Kaiyu Zheng, Leonid P. Savtchenko, Dmitri A. Rusakov

**Affiliations:** UCL Queen Square Institute of Neurology, University College London, United Kingdom; Neuroscience and Cell Biology Research Institute, City St George’s University of London, Cranmer Terrace, London SW17 0RE, United Kingdom

**Author notes:** Equal contribution (Olga Kopach, James P. Reynolds).

## Abstract

Information handling and storage by neural circuits is thought to involve dynamic changes in the synaptic connectome, yet how this process unfolds at the level of individual synapses in vivo remains unclear. We took advantage of a well-defined thalamocortical anatomical framework to monitor identifiable individual synapses in barrel cortex during rhythmic whisker stimulation (RWS). Multiplexed ratiometric readouts from genetically targeted optical sensors showed that RWS-induced long-term potentiation (LTP) increased glutamate release per action potential in RWS-responsive axons while recruiting previously silent thalamocortical connections. Fast high-resolution imaging revealed pronounced inter-synaptic glutamate transients and global extracellular GABA waves triggered by brief RWS. LTP induction had no detectable effect on the evoked GABA signal but further enhanced glutamate crosstalk beyond thalamocortical synapses. Strikingly, neural-network simulations suggest that such volume-transmitted excitatory signals can improve associative memory retrieval in sparsely connected networks. Together, these findings uncover key plasticity features of the cortical synaptic connectome and point to a potential computational consequence of glutamate spillover for brain circuit function.

## INTRODUCTION

Excitatory glutamatergic synapses are fundamental to brain function. Their ability to modify transmission strength in a coincidence-dependent manner forms the basis of Hebbian learning principles ^1^. These principles have found a robust experimental counterpart in synaptic long-term potentiation (LTP) ^2^, a phenomenon now closely linked to memory formation ^3–7^. Yet two fundamental questions remain unresolved. First, how is synaptic potentiation expressed at individual identified synapses in the intact brain: through presynaptic changes, postsynaptic changes, or both? Second, how can cortical circuits enhance information processing without simply increasing excitation? Proposed stabilizing mechanisms include homeostatic synaptic scaling ^8–10^, alterations in intrinsic neuronal excitability ^11–13^, or a combination of both ^14–16^. Addressing both questions has proved difficult because distinguishing changes in synaptic efficacy from alterations in cellular or network excitability in vivo remains technically challenging. Likewise, recent demonstrations of memory-associated changes in synaptic connectivity ^17–19^ raise the further question of whether circuit plasticity is mediated primarily by changes in postsynaptic responsiveness, presynaptic release probability, or dynamic redistribution of functional connectivity within existing networks.

In this context, a well-established theory holds that optimal information handling in cortical networks relies predominantly on control of release probability P_r_ ^20–23^. Indeed, P_r_ at small excitatory synapses can vary more than fivefold ^24–27^, offering considerable potential for activity-dependent tuning of synaptic connectivity including LTP induction ^28–30^. Yet, establishing the roles of P_r_ dynamics in excitatory circuit plasticity *in vivo*, under physiological stimulation, has remained a significant challenge.

This challenge extends to a long-standing question of whether individual glutamatergic synapses provide point-precision, one-to-one signaling devices, or whether excitatory signals can spread to neighboring connections. Although activation of only several dozen postsynaptic receptors is sufficient for synaptic transmission, thousands of glutamate molecules released into the synaptic cleft escape into the surrounding neuropil, where they are rapidly captured by high-affinity transporters ^31,32^, predominantly the astrocytic transporter GLT-1 ^33,34^. Nevertheless, electrophysiological studies in vitro and in brain slices have long suggested that escaping glutamate can potentially activate high-affinity receptors up to several microns from the release site ^35–38^. More recently, experiments employing genetically encoded glutamate sensors have corroborated these observations ^25,39,40^. Because such glutamate “spillover” challenges the traditional view of neural circuits as collections of discrete one-to-one synaptic connections, the extent to which it occurs in the intact brain remains a major unresolved question.

Here, we sought to address these challenges by monitoring glutamate release and presynaptic Ca^2+^ dynamics at individual thalamocortical synapses in the barrel cortex, under the experimental paradigm of rhythmic whisker stimulation (RWS) ^41^ (Figure 1A), including RWS-induced LTP of the sensory signal ^42–44^. We therefore focused on the circuitry of the posterior medial nucleus (POm) of the thalamus, which sends its axonal projections to the barrel cortex in S1 (mainly layer L1) ^45–48^ where ramifications are widespread ^49,50^. While this thalamocortical circuitry is itself the subject of intense ongoing research ^43–47,49–51^, here it provides a well-defined anatomical framework that allows us to isolate and interrogate individual cortical synapses of known origin.

**Figure 1.**
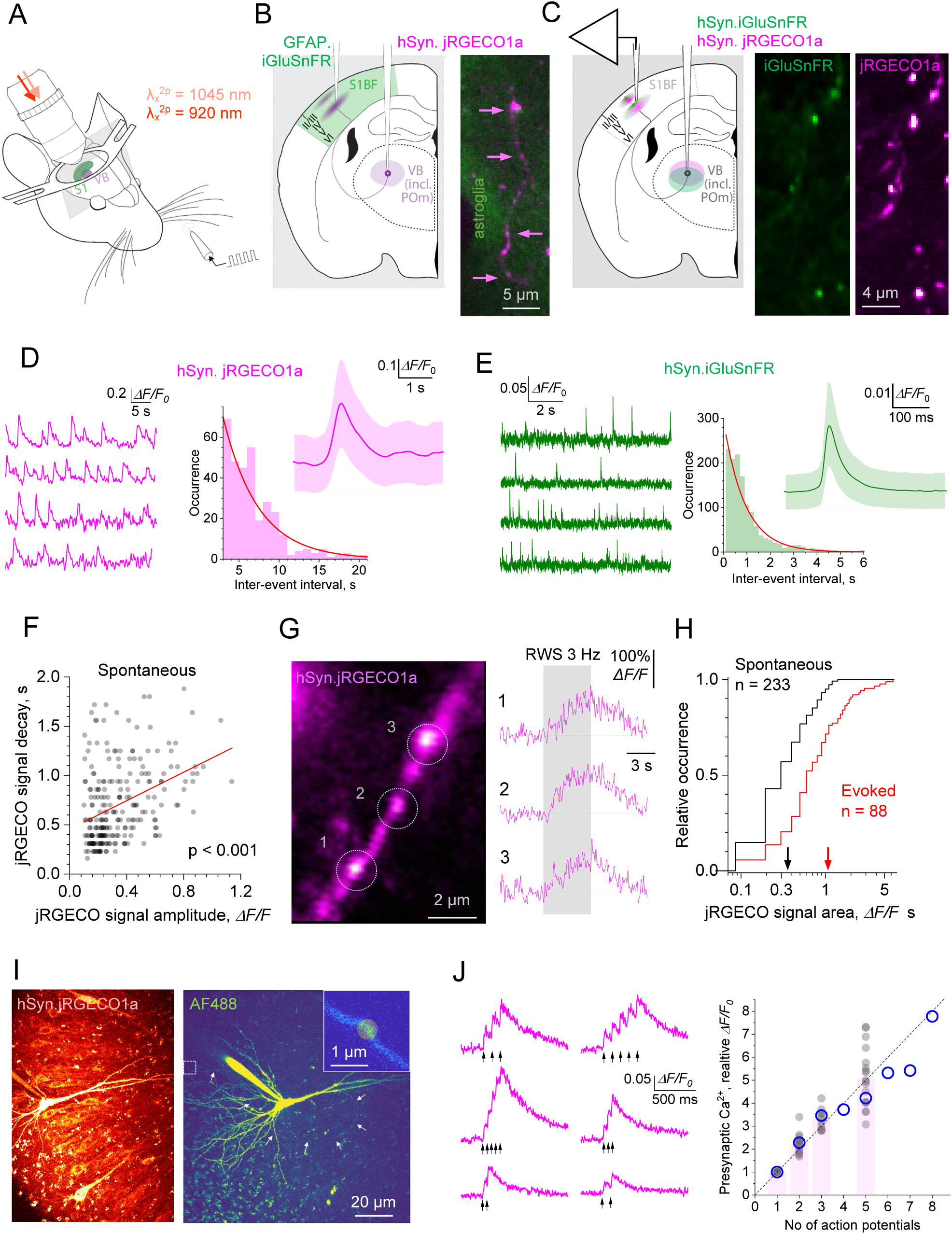
Multiplexed imaging of glutamate release and presynaptic Ca^2+^ at individual thalamocortical synapses. **(A)** Experimental arrangement: RWS setting with a cranial window over the contralateral barrel cortex for two-photon excitation microscopy. **(B)** Diagram: viral transduction of hSyn.jRGECO1a and GFAP.iGluSnFR.A184S in thalamic POm neurons and S1BF astroglia, respectively; image: cranial window snapshot depicting expression of jRGECO1a in thalamocortical axons (magenta, arrowheads) and iGluSnFR in local astroglia (green). **(C)** Diagram: dual-viral transduction of hSyn.jRGECO1a and hSyn.iGluSnFR .A184V in POm neurons, with electrophysiological control of RWS in S1BF. **(D)** Traces: examples of spontaneous Ca^2+^ events (jRGECO1a channel) in individual boutons of POm axons, sensor expression as in **B**; plot, histogram of inter-event intervals (mean ± SEM: 700 ± 33 ms; n = 384 events, 6 boutons; analysis example in Figure S2A); inset, average event shape (mean ± SEM). **(E)** Traces: examples of spontaneous glutamate release events (iGluSnFR channel) in individual boutons of POm axons, sensor expression labelling as in **C**; plot, histogram of inter-event intervals (mean ± SEM: 17.3 ± 0.2 ms; n = 1342 events, 5 boutons; analysis example in Figure S2C); inset, average shape of spontaneous events (± SEM). **(F)** Uniformity test for axonal Ca^2+^ signals: decay constant of spontaneous presynaptic Ca^2+^ events recorded as in **B**, plotted against the *ΔF/F0* signal amplitude (jRGECO1a channel, n = 233 events); line, linear regression (slope at p < 0.001). **(G)** Image: example of a POm axonal fragment with three jRGECO1a-expressing boutons (1-3, left); traces: fluorescence responses evoked by RWS (5 s at 3 Hz; grey segment, right). **(H)** Frequency distribution of the jRGECO1a signal area under *ΔF/F_0_* curve (AUC), for spontaneous and evoked events at POm boutons, as in **D** and **G**. **(I)** Experiments to test relationship between spike number and presynaptic jRGECO1a signal: *left*, CA3 pyramidal cell expressing jRGECO1a held whole-cell; *right*, axon traced using dialyzed AF488, with a selected synaptic bouton imaged using a Tornado linescan (inset, spiral). **(J)** Linearity test: traces, examples of presynaptic jRGECO1a responses evoked by varied spike bursts; plot, jRGECO1a response plotted against the spike total (grey dots, individual experiments; columns, mean; blue circles, data from mouse visual cortex ^52^; dotted line, 1:1 linear dependence).

## RESULTS

### Multiplexed imaging of individual thalamocortical axons under whisker stimulation

To monitor activity of individual thalamocortical connections, we used two approaches. In the first approach, we virally transduced the red Ca^2+^ indicator jRGECO1a ^52^ in POm neurons, and the green glutamate sensor iGluSnFR ^53,54^ in S1BF area astrocytes (Figure 1B, Figure S1A), using procedures established previously ^39,48^ (STAR Methods). Astrocytic expression of iGluSnFR was chosen to enable detection of glutamate signals beyond the boundaries of targeted POm axons. Two-three weeks after AAV transduction, a cranial window was implanted over S1BF, so that both glutamate and Ca^2+^ signals induced by RWS at POm synapses, could be detected using wide field-scanning mode (Figure S1B) and investigated further at higher spatiotemporal resolution. In the second approach, we expressed both sensors in POm neurons (STAR Methods), to restrict both jRGECO1a and iGluSnFR signaling to the cortical synapses that successfully expressed both sensors (Figure 1C).

In rodent barrel cortex layer L1, the volume density of POm-projecting synapses is ∼0.047 μm^-3^ (Refs ^55,56^). Assuming an approximately 1 µm-thick two-photon excitation focal plane, the average volume density of jRGECO1a-labelled boutons, estimated from their area densities across experimental ROIs, was 0.0058 ± 0.0013 µm^-3^ (mean ± SEM, n = 25 ROIs in 25 animals; total 809 boutons registered for this purpose). This suggests that jRGECO expression was detected in approximately 12% of POm axons. In awake animals, 20-30% of the targeted axons showed detectable spontaneous events (Figure 1D-E, traces). The analyses (Figure S2A, C) revealed that the event sequences were indistinguishable from the Poisson process in either jRGECO1a or iGluSnFR channel (Figure 1D-E, plots). A positive association between event amplitude and its decay and frequency (Figure 1F, Figure S2B, D) suggested non-unitary, compound spike-burst events.

Introducing light anesthesia reduced spontaneous activity severalfold, enabling detection of RWS-evoked presynaptic Ca²⁺ responses which were 2-5 times longer than spontaneous events, suggesting longer spike bursts (Figure 1G). The evoked glutamate signal varied considerably more than presynaptic Ca^2+^ signal (Figure S2E-G; here ’failures’ report signals indistinguishable from noise), consistent with the probabilistic nature of glutamate release ^25^. Because direct estimation of synaptic P_r_ was not feasible under these experimental conditions, we instead sought to quantify synaptic release efficacy (E_r_) defined as the relative amount of glutamate released per action potential (or spike-evoked Ca^2+^ entry) during a 200-ms RWS (four stimuli at 20 Hz).

While the iGluSnFR signal typically scales with the amount of glutamate released during short spike bursts ^25,53,54,57–59^, estimating E_r_ also requires knowing the number of presynaptic action potentials during each 200-ms RWS trial. The magnitude of non-saturating somatic Ca²⁺ signals has been shown to exhibit a near-linear relationship with spike number ^52,60,61^, which prompted us to examine this relationship for presynaptic jRGECO1a under our experimental conditions. First, we confirmed that presynaptic jRGECO1a responses evoked by relatively long (3 s) RWS (Figure 1G) remained well below saturation (Figure 1H). Second, experiments in brain slices confirmed a linear relationship between spike number and presynaptic jRGECO1a signal (AUC) across a range of spike bursts (Figure 1I-J), consistent with previous observations in mouse visual cortex ^52^. Together, these results indicate that the ratio between iGluSnFR and jRGECO1a responses to brief RWS (hereafter referred to as Glu/Ca) scales with the average amount of glutamate released per spike-evoked Ca^2+^ entry, or release efficacy E_r_. In an additional test, we showed a direct relationship between Glu/Ca ratio and P_r_ using another red-shifted Ca^2+^ indicator (Figure S3).

### RWS-induced LTP boosts glutamate release efficacy at thalamocortical synapses

We confirmed that the established LTP induction protocol in this circuitry (RWS at 3 Hz for 120s) ^42–44^ produced robust potentiation of field EPSPs induced by 200-ms RWS in the barrel cortex, lasting at least 60 min (Figure 2A-C). We therefore were able to record synaptic responses evoked by 200-ms RWS, in baseline conditions and 10-50 min following RWS-LTP induction, using high-speed (0.5-1kHz) Tornado scans, as detailed previously ^25^, under both astrocytic and axonal expression of iGluSnFR (Figure 2D-H).

**Figure 2.**
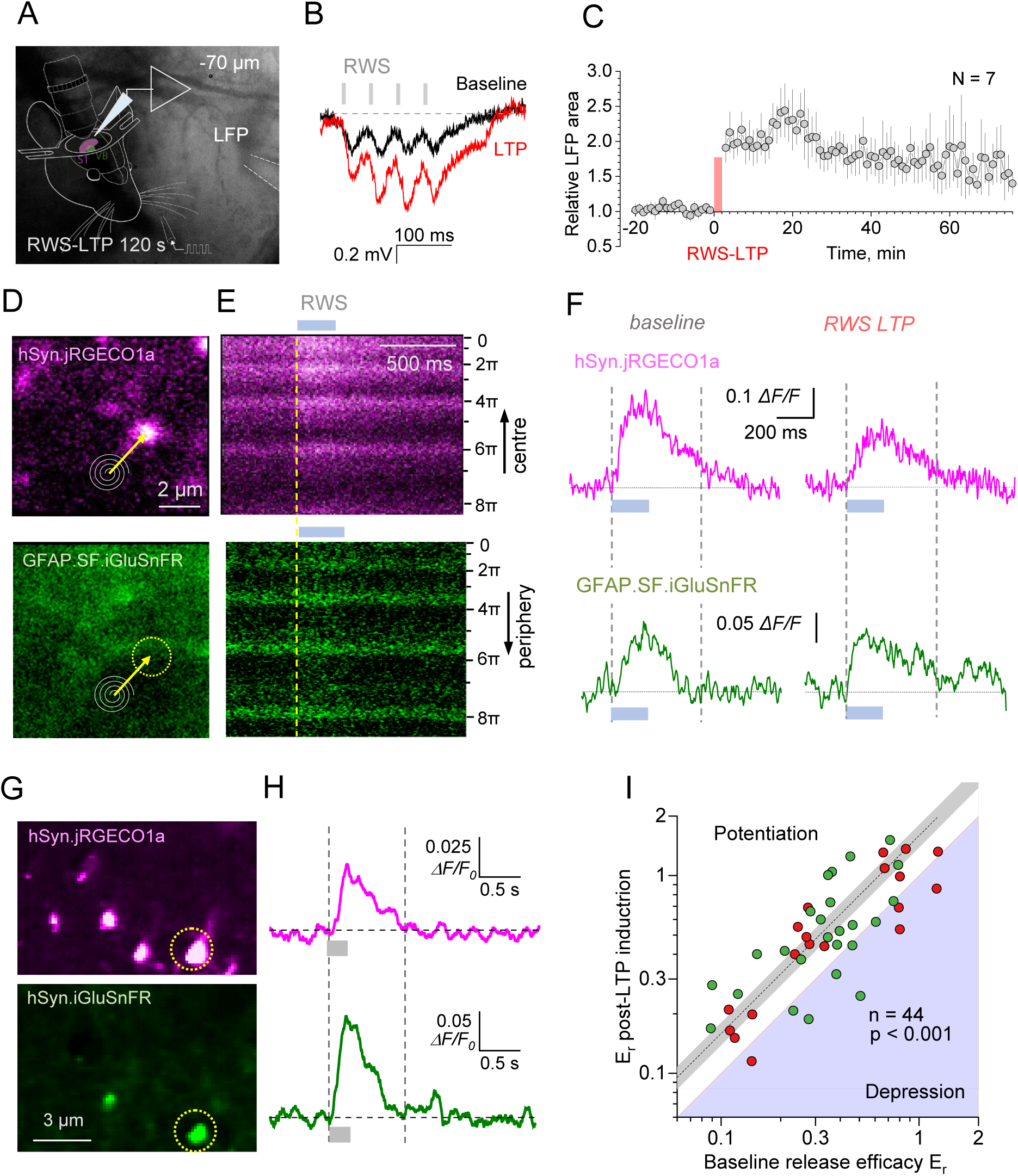
RWS-induced LTP increases glutamate release efficacy and recuites additional thalamocortical connections. **(A)** Experimental arrangement: electrophysiological control of RWS-induced LTP. **(B)** Barrel cortex fEPSPs evoked by 200-ms RWS (four puffs at 20 Hz) in baseline conditions (black) and 20 min after RWS-LTP protocol (red, 120 s at 3 Hz), as indicated. **(C)** fEPSP magnitude (AUC, mean ± SEM, n = 7 animals) before and after RWS-LTP induction (red segment). **(D)** Multiplexed imaging of a jRGECO1a-expressing POm axonal bouton (top, magenta) and local iGluSnFR-expressing astroglia (bottom, green); dotted circle, Tornado scan ROI; spirals and arrows, Tornado scan position. **(E)** Tornado linescan (∼500 Hz) of the axonal bouton in **d**, showing a fluorescent transient upon 200-ms RWS (grey segment; dotted line, onset) in both channels; ordinate, rotation angle of the Tornado spiral (radians). **(F)** ROI-integrated Tornado-scan fluorescence time course in jRGECO1a and iGluSnFR channels, as indicated, during 200-ms RWS (grey segment), before (left) and 10-30 min after RWS-LTP induction (right, average of four trials); dotted lines, sampling windows to calculate the Glu/Ca ratio (using *ΔF/F_0_* AUC). **(G)** Multiplexed imaging of POm boutons expressing both jRGECO1a (top) and iGluSnFR (bottom); dotted circle, bouton of interest ROI. **(H)** ROI-integrated Tornado-scan fluorescence time course in jRGECO1a and iGluSnFR channels, as indicated, during 200-ms RWS (grey segment), 15-1 min before (average of 8 trials, left) and 5-35 min after RWS-LTP induction (average of 10 trials, right); other notations as in **F**. **(I)** Summary, changes of glutamate release efficacy E_r_ during LTP: Glu/Ca ratio after LTP induction plotted against Glu/Ca ratio in baseline conditions for 44 individual POm synapses; red and green circles, data under the labelling protocols shown in **D-F** and **H-G**, respectively; dotted line ± shaded area, mean change ± SEM (p < 0.001, n = 44 in 16 animals). **(J)** Changes in E_r_ plotted against the baseline E_r_ for individual synapses; dotted line, linear regression (p < 0.004, n = 44, *t*-test). **(K)** Relative change in *ΔF/F_0_* iGluSnFR and jRGECO1a responses 20-30 min after RWS-LTP induction; * p < 0.03 (single-sample *t*-test), ***, p < 0.001 (paired-sample *t*-test), n = 44. **(L)** Image, snapshot of two jRGECO1a-expressing POm axonal boutons, 1 and 2, in SB1 area; traces, Ca^2+^ response (jRGECO1a channel) of boutons 1 and 2 to 200-ms RWS (grey segment) before (grey trace) and 20-30 min after (red) RWS-LTP induction, as indicated. **(M)** Summary of Ca^2+^ activity at individual POm axonal boutons (n = 285, 13 animals), indicating initially 216 silent (grey), 49 initially responsive (magenta), and 20 boutons responsive only after LTP induction (red); see Figure S5 for further example and detail.

In individual boutons, LTP induction was followed by a significant increase in synaptic release efficacy, E_r_: Glu/Ca ratio increased by 59 ± 10% (mean ± SEM; p < 0.001; n = 44 boutons from 16 animals; Figure 2I, data color-coded according to the two dual-sensor expression protocols). This effect could not be explained by divergent photobleaching trends of the two sensors (Figure 3A). Furthermore, baseline E_r_ values were negatively correlated with their post-LTP change (Figure 3B), in line with previously reported behaviour of P_r_ inferred from optical quantal analysis during LTP induction ^28^.

**Figure 3.**
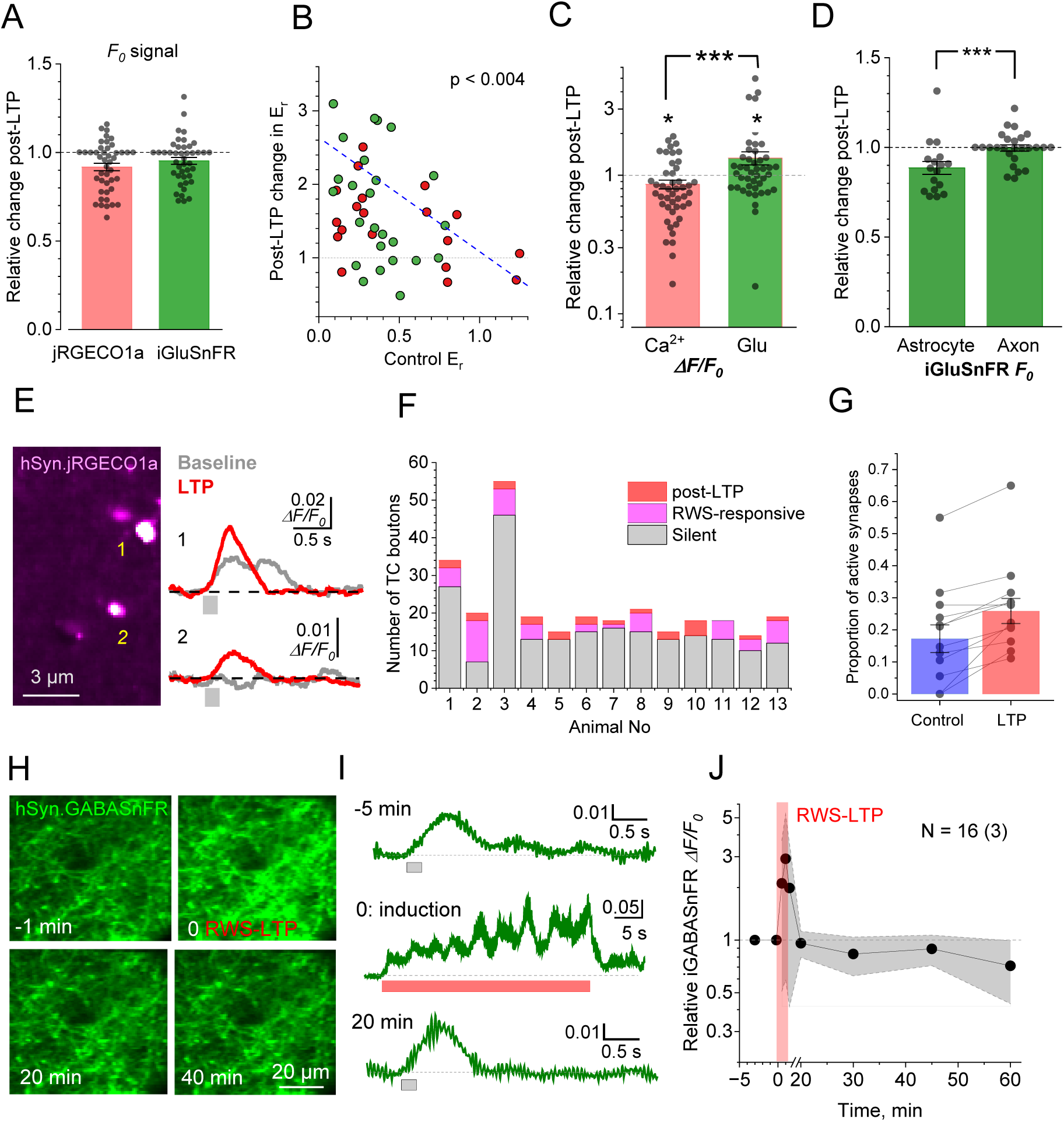
LTP induction increases glutamate release at active and recruits additional POm synapses but leaves overall inhibitory signal in L1 unchanged. **(A)** Photobleaching test: Relative changes in baseline fluorescence (*F_0_*) recorded at presynaptic boutons of thalamocortical axons in jRGECO1a (−8.2 ± 2.1%) and iGluSnFR (−4.8 ± 2.0%; astrocytic and axonal expression combined) channels, as indicated, after the induction of RWS-LTP; dots, individual bouton data (bars, mean ± SEM; n = 43 boutons in 16 animals). **(B)** Changes in glutamate release efficacy E_r_ at individual POm synapses plotted against the efficacy in control conditions; dotted line, linear regression (p < 0.004, n = 44, regression *t*-test). **(C)** Relative change in *ΔF/F_0_* signals of glutamate (iGluSnFR channel) and Ca^2+^ (jRGECO1a channel) 20-30 min after RWS-LTP induction; * p < 0.03 (single-sample *t*-test), ***, p < 0.001 (paired-sample *t*-test), n = 44. **(D)** Relative changes in the baseline fluorescence (*F_0_*) of astrocyte-expressed iGluSnFR (integrated over the focal plane within the Tornado ROI, as in Fig. 3D, bottom; −11.5 ± 3.6%, n = 17) and of POm axon-expressed iGluSnFR (−0.1 ± 1.8%, n = 26); ***, p < 0.005. **(E)** Image, snapshot of two jRGECO1a-expressing TC axonal boutons, 1 and 2, in SB1 area; traces, Ca^2+^ response (jRGECO1a channel) of boutons 1 and 2 to a test RWS stimulus (200 ms at 40 Hz, grey segment) before (grey trace) and 20-30 min after (red) RWS-LTP induction, as indicated. **(F)** All recorded individual POm axonal boutons (285 boutons, 13 animals), including Ca^2+^-silent ones, before and after RWS-LTP induction (see Figure S5 for further example). **(G)** Summary of experiments shown in **E-F**: proportion of RWS-responsive POm axonal boutons in baseline conditions and after LTP induction; ***, p < 0.001 (n = 13 animals; data combined for both protocols of dual sensor expression). **(H)** Frame-scan snapshots of iGABASnFR2-expressing neuronal processes in SB1 (depth ∼100 µm) before (−1 min), during (0 min), 20 min and 40 min after the RWS-LTP induction protocol, as indicated. **(I)** Characteristic frame-integrated fluorescence responses (iGABASnFR2 channel, resonant frame scanning, 1 kHz) to RWS (200 ms at 40 Hz, grey segment) in baseline conditions (top trace), during RWS-LTP induction protocol (middle), and 20 min after the induction (bottom); single-trial traces. **(J)** Summary of experiments shown in **H-I**: relative magnitude (AUC, mean ± 95%CI) of iGABASnFR2 responses to test RWS (200 ms at 40 Hz) before, during, and after the RWS-LTP induction (red segment), as indicated; n = 16 brain areas / 3 animals.

The LTP-induced increase in E_r_ appeared to reflect both an increase in the RWS-evoked iGluSnFR signal post-LTP (33 ± 14%, p < 0.05, n = 44) and, perhaps surprisingly, a decrease in the presynaptic jRGECO1a signal (14 ± 6%, p < 0.05; Figure 3C). The latter may reflect either a reduction in the number of RWS-evoked burst spikes or, albeit less likely ^30^, a decrease in action potential-driven Ca^2+^ entry. Under these conditions, LTP induction had no detectable effect on either the frequency or amplitude of spontaneous glutamate release events (Figure S4C-D).

The baseline fluorescence signal (*F_0_*) generated by iGluSnFR-expressing astroglia and collected during bouton-centered Tornado scans (across an approximately 1 µm-thick two-photon excitation focal plane) scales with the tissue volume fraction occupied by perisynaptic astroglia ^39,62^. Intriguingly, this signal decreased during RWS-LTP by 11.5 ± 3.6% (p < 0.01, n = 17), whereas the *F_0_* signal from iGluSnFR-expressing axons remained remarkably stable (change: −0.1 ± 1.8%, n = 26; Figure 3D). The latter observation rules out the effect of photobleaching on astrocyte-expressed iGluSnFR and suggests a reduced presence, or partial withdrawal of perisynaptic astroglia near potentiated synapses, consistent with our earlier findings ^39^ and a recent 3D electron microscopy study^63^.

### RWS-induced LTP recruits additional thalamocortical connections

Nonetheless, the 33% net increase in glutamate signal following LTP induction (Figure 3C) could not fully explain the 80-100% potentiation level reported by electrophysiology (Figure 2C). We therefore hypothesized that RWS-LTP, while reducing excitation in the originally responsive pathway, could recruit neuronal projections that were previously silent. Going back to the original image series we found that out of 285 registered axonal boutons (across 13 animals), 49 were active at baseline, compared to 72 after LTP induction (Figure 3E-F, Figure S5), a 47% increase. This represents a significant change in the proportion of active POm synapses (from 17 ± 4% to 25 ± 4%, mean ± SEM, n = 13 animals, p < 0.001; Figure 3G). Together with a 33% increase in overall glutamate release, this change is consistent with the expression level of LTP documented by electrophysiology. These findings provide evidence for a dynamic (re)allocation of thalamocortical pathways, consistent with hypotheses previously suggested by current-source density mapping ^42^ and by monitoring local network activity ^64^ during RWS-induced plasticity.

### Overall inhibitory signal remains unchanged after RWS-induced LTP

An important mechanism that could influence the efficacy of thalamocortical synapses during RWS-LTP involves the extensive inhibitory circuitry of barrel cortex L1. Local interneurons play a key role in shaping sensory maps through their interactions with sensory activity ^65–67^ and contribute critically to RWS-LTP in layer 2/3 principal neurons ^64,68^. While interneuron firing patterns recorded with Ca^2+^ imaging provides a powerful readout of local network activity, the transmitted inhibitory signal ultimately depends on the spatiotemporal dynamics of GABA release. Unlike glutamate, synaptically released GABA encounters no high affinity transporters and can therefore diffuse over tens of microns, generating substantial extracellular waves ^69–71^. The absence of rapid transporter-mediated buffering provides an excellent opportunity to monitor GABA release patterns using recently developed optical GABA sensors ^70,72^.

To assess the overall contribution of GABA signaling to RWS-LTP in POm synapses, we monitored extracellular GABA dynamics using the genetically encoded sensor iGABASnFR2 ^70^, expressed broadly in cortical neuronal processes (Figure S6A-B; STAR Methods). A 200-ms RWS induced robust iGABASnFR2 responses that were consistently detectable over large spatial scales, albeit with some local heterogeneity (Figure S6C). The RWS-LTP induction sequence triggered a several-fold increase in GABA signal (Figure 3H-I), yet iGABASnFR2 responses to 200-ms RWS remained unchanged thereafter (Figure 3J, n = 16 ROIs in 3 animals). Although these data cannot address possible LTP-associated re-arrangements of active interneuronal circuits ^64,68^, they argue against a major change in overall local inhibitory signaling in the maintenance of RWS-LTP at POm synapses.

### Brief RWS triggers substantial glutamate spillover from thalamocortical synapses

Extrasynaptic glutamate escape has been extensively documented in brain slice preparation ^25,36,38–40^. However, its extent in the intact brain remains largely unknown, and its functional significance is still poorly understood.

To address this, we used high-resolution 2D reconstructions of Tornado scans ^25^ to relate profiles of jRGECO1a-filled axonal boutons (Figure 4A, top) to the fluorescence landscapes of iGluSnFR expressed in the surrounding astroglia (Figure 4A, bottom). The numerical ratio between iGluSnFR images over a 200-ms period during and 200 ms before the RWS-induced response (Figure 4B) provided a *ΔF/F_0_* signal landscape reporting a projected profile of glutamate escape in the synaptic proximity (Figure 4C, top). In most cases, the *ΔF/F_0_* image revealed a hotspot, pointing to the tentative glutamate release site, with the signal fading with distance (Figure 4C; see Figure S7 for additional examples).

**Figure 4.**
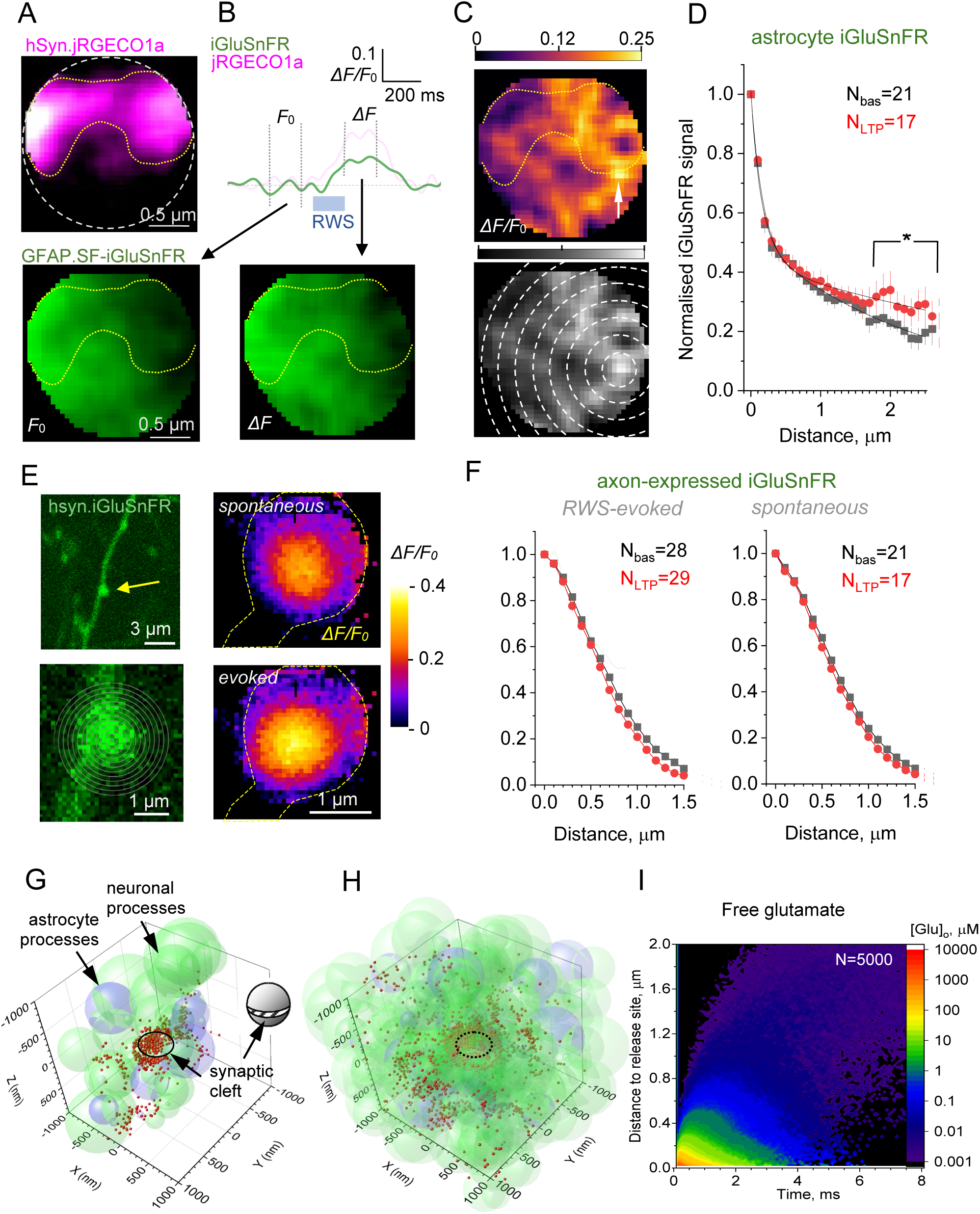
RWS-induced glutamate spillover at thalamocortical synapses. **(A)** 2D-reconstructed image of high-resolution Tornado scan (500 Hz, 200-ms average), showing a jRGECO1a-expressing POm axonal bouton (top) and the iGluSnFR-expressing surrounding astroglia (bottom), in resting conditions; dotted white circle, Tornado scan area; yellow dotted line, axonal bouton outline, in accordance with its cytosolic jRGECO1a fluorescence. **(B)** Traces, area-integrated Tornado-scan fluorescence time course in iGluSnFR and jRGECO1a channels, as indicated, during 200-ms RWS (grey segment); vertical dotted lines, time windows to average 2D landscapes of iGluSnFR signal; image (bottom), 2D landscape of iGluSnFR fluorescence as in **A**, but during 200-ms RWS (*ΔF* sampling window indicated). **(C)** Top, *ΔF/F_0_* iGluSnFR signal landscape: the ratio of *ΔF* image in **B** and *F*_0_ image in **A**; bar, false-color signal scale; arrow, hotspot (fluorescence peak). Bottom, image as above, but in brightness-calibrated grey scale, with concentric circles to calculate average signal decay from the hotspot; see Figure S7 for further examples. **(D)** Spatial decay profile of RWS-evoked astroglial *ΔF/F_0_* iGluSnFR signal, normalized with respect to hotspot fluorescence, in control conditions (grey squares, n = 21) and 10-30 min after RWS-LTP induction (red circles, n = 17; mean ± SEM); solid lines, best-fit biexponential approximation in the form *A_1_*·exp(−x/λ_1_) + *A*_2_·exp(−x/λ _2_); control best-fit shown: *A*_1_ = 0.491, λ_1_ = 0.134 μm, *A*_2_ = 0.518, λ_2_ = 2.38 μm; LTP best-fit: *A*_1_ = 0.572, λ_1_ = 0.181 μm, *A*_2_ = 0.431, λ_2_ = 5.42 μm; the average long-range (>1.5 µm) decay profile elevates after RWS-LTP induction by 58 ± 26%, as indicated; *, p < 0.05 (two-sample *t*-test; see Figure S8 for further detail); N, bouton number in baseline and LTP conditions, as indicated (4 boutons were lost post-induction). **(E)** iGluSnFR-expressing POm axon (top left) with a presynaptic bouton (arrow; shown at high magnification with a Tornado scan position, bottom) displaying *ΔF/F_0_* iGluSnFR signal landscape (average of 5-7 events, 15 ms average over the peak signal) during spontaneous (top right) and RWS-evoked (bottom right) glutamate events; dotted line, outer optical boundary of the bouton; other notations as in **C**. **(F)** Analyses as in **D**, but for the Pom axon-expressed hSyn.iGluSnFR during RWS-evoked (left) and spontaneous (right) events; grey shade, previously reported spatial decay of hSyn.iGluSnFR signals (mean ± 95CI, n = 24 axons) in CA3 cell axonal boutons, hippocampal slices, adapted from ^25^; other notations as in **D**. **(G)** Snapshot of Monte Carlo simulations (500 nm wide sample of the 4 x 4 x 4 µm^3^ arena) showing glutamate molecules (red dots) 4 ms after release inside the synaptic cleft (see Figure S9A), with a stochastically generated perisynaptic neuropil containing neuronal (green) and astrocyte (pale blue-violet) processes represented by overlapped concatenated spheroids. Model parameters represent layer 1 cortical neuropil ^73,74^: ∼20% volume fraction for extracellular space, ∼10% astroglia, ∼70% neuronal elements; see Methods for further detail ^75^. At 4 ms post-release, >90% glutamate molecules are bound to transporter-enriched astrocyte surfaces. **(H)** Snapshot in **G** expanded to a 2 x 2 x 2 µm^3^ arena (see Figure S9C for the same snapshot with astroglia only shown). **(I)** Example of the spatiotemporal dynamics of free (non-bound) glutamate in simulation experiments shown in **G-H**, with glutamate binding sites expressed throughout astroglial surfaces at Ψ = 0.1 ms.

Analysis of the signal decay relative to the hotspot in such images (Figure 4C, bottom) suggested that the RWS-evoked glutamate transient extends at least ∼2.5 μm from the release site (Figure 4D). The volume densities of POm-projecting and all excitatory synapses in barrel cortex L1 are ∼0.047 μm⁻³ and ∼0.79 μm⁻³, respectively ^55,56^, corresponding to average nearest-neighbor distances of ∼1.5 μm and ∼0.6 μm ^56^. Given that ∼20% (17-25%) of POm axons responded to RWS (Figure 3G), the nearest-neighbor distance between RWS-activated POm synapses in our experiments was 1.5 · 0.2^−1/3^ ∼ 2.6 µm. Thus, the *ΔF/F₀* iGluSnFR decay profile (Figure 3G) is likely to reflect overlap between glutamate escape profiles originating from both POm synapses and neighboring excitatory synapses in the area. Thes observations point to the possibility of inter-synaptic communication mediated by glutamate receptors, at least those with affinities comparable to iGluSnFR (see below).

### RWS-LTP increases inter-synaptic but not perisynaptic glutamate signal

We have previously shown that RWS-LTP in this circuitry induces the withdrawal of perisynaptic astroglia from potentiated synapses ^39^, which is consistent with present observations (Figure 3D). Because astrocytes are enriched in high affinity glutamate transporters ^39^, such changes suggested enhanced extrasynaptic glutamate transient. Indeed, RWS-LTP elevated the extrasynaptic glutamate profile, but only at distances >1.5 μm from the hotspot of POm boutons (by 48%; Figure 4D, Figure S8). As discussed above, this is likely to reflect greater overlap among glutamate transients generated by neighboring synapses, rather than an increase in the glutamate signal immediately adjacent to individual potentiated synapses. Such enhanced overlap is also consistent with the LTP-induced recruitment of previously silent POm connections, which increased the number and hence the volume density of RWS-responsive thalamocortical synapses in our experiments (Figure 3F-G).

To examine this further, we analyzed recordings in which both sensors were expressed exclusively in axonal boutons (Figures 1C, 2G-H), thereby reporting glutamate dynamics in the immediate vicinity of the release site (Figure 4E). This analysis revealed a markedly more restricted glutamate escape profile for both evoked and spontaneous release events (Figure 4F) compared with astrocyte-expressed iGluSnFR (Figure 4D). This difference is consistent with the absence of astrocytic iGluSnFR signal integration across the surrounding tissue volume when the sensor is confined to individual presynaptic boutons. Under these conditions, the apparent extent of glutamate spillover closely matched the previously reported *ΔF/F_0_* iGluSnFR profile at individually activated synapses in slices (Figure 4F, grey shading) and showed little change following RWS-LTP induction (Figure 4F). These observations further support the notion that the enhanced extrasynaptic glutamate signal detected after RWS-LTP reflects increased overlap among glutamate transients originating from neighboring synapses rather than greater glutamate escape from individual potentiated synapses.

### Could escaping glutamate activate its receptors at neighboring synapses?

These observations suggest that glutamate discharged in response to brief RWS synapses generates significant transients outside the POm projections. However, as iGluSnFR reports only sensor-bound glutamate, the fate of free glutamate molecules has remained uncertain ^76^. To investigate this scenario through detailed biophysical modelling, we generated a realistic perisynaptic neuropil built on our previous simulations of the microenvironment of cortical synapses ^77–79^. In this model, a 300 nm wide, 20 nm high synaptic cleft is enclosed between two hemispheres representing persistent pre-and postsynaptic obstacles to diffusion ^77,80^ and surrounded by stochastically generated neuronal and astrocyte elements ^75^ (Figure 4G-H, Figure S9A), occupying 10% and 70% tissue volume fraction, respectively, values typical for barrel cortex L1 ^73,74^, leaving ∼20% for the extracellular space ^81,82^. In these simulations, sub-microscopic cellular processes are represented by overlapping spheroids of varied diameters and positions (to make caterpillar-like, intersecting structures), reflecting the arbitrary nature of cell morphology on the nanoscale. We validated our diffusion algorithms by comparing the outcome of our Monte Carlo simulations with the theoretical predictions of Maxwell’s diffusivity for a sphere-filled porous medium with similar parameters (Figure S9B).

The astrocytic membranes expressing high-affinity iGluSnFR in our glutamate spillover trials (Figure 4E) are also densely populated with high-affinity GLT-1 glutamate transporters, which buffer extracellular glutamate under native conditions ^31,33,83^. Both the sensor and the transporter have very high glutamate association rates, on the order of 10^6-^10^7^ M^-1^s^-1^ (ref. ^84^), making extrasynaptic glutamate buffering immediately after release effectively diffusion-limited, i.e. quasi-instantaneous locally. Given their shared astrocyte localization and glutamate-binding properties, the combined concentration of GLT-1 and iGluSnFR must shape the extrasynaptic glutamate profile reported by iGluSnFR fluorescence decay relative to the hotspot ^76^.

Because the exact extracellular concentrations of iGluSnFR and GLT-1 are difficult to evaluate, we simulated 3D decay profiles of bound glutamate following single-synapse release of 5000 glutamate molecules over a 10^6^-fold buffer concentration range represented by parameter Ψ (Figure S9D, black lines; all other modelling parameters were constrained by experimental data; STAR Methods). These distributions were compared with the 3D profile of RWS-evoked iGluSnFR signal profiles, reconstructed from recorded 2D projections (Figure 4D) by Abel Inversion (STAR Methods), for both astrocyte-and POm axon-expressed sensors (Figure S9D). The best-fitting profile of the axon-expressed iGluSnFR signal corresponded to Ψ ≈ 0.1 ms (Figure S9D). This enabled us to estimate the spatiotemporal dynamics of unbuffered glutamate (Figure 4I) and its potential action at NMDA receptors, key high-affinity receptors expressed at excitatory synapses (Figure S9E).

It is important to emphasize that the present estimates are conservative. Glutamate buffering in our experiments is augmented by the presence of iGluSnFR ^76^, while the simulations do not account for glutamate unbinding from GLT-1 and iGluSnFR beyond ∼8-10 ms after release ^75^ or for multivesicular release. Each of these factors would be expected to increase glutamate escape and the probability of extrasynaptic receptor activation.

Together, these analyses support the notion that RWS triggers excitatory inter-synaptic crosstalk beyond POm axons, likely engaging multiple local glutamate sources ^55,56^ and exposing high-affinity receptors at nearby inactive synapses to glutamate concentrations sufficient for receptor binding and potentially activation.

### Potential network consequences of glutamate-mediated inter-synaptic crosstalk

The role of glutamate spillover in shaping cortical network performance remains poorly understood, and in cortical L1 the interaction between long-range inputs and local circuitry is complex. Learning-related plasticity occurs in the local branches of pyramidal neurons, with synaptic changes in this layer reflecting sensorimotor learning ^85,86^. However, the blurring of one-to-one synaptic connectivity by escaping glutamate has not traditionally been viewed as compatible with accurate information transfer or storage.

To test how excitatory inter-synaptic crosstalk can affect these basic operational features of a neural network, we examined associative memory recall in Hopfield networks consisting of spiking neurons ^87,88^. The networks were trained to store three overlapping memory patterns, one of which was subsequently presented as a probe (cue), either noise-free or contaminated by varied levels of noise (Figure 5A). The network recall quality *Q* (STAR Methods) was assessed across 10 independent trials for each probe and condition (network performance at *Q* < 40% was judged poor and therefore ignored).

**Figure 5.**
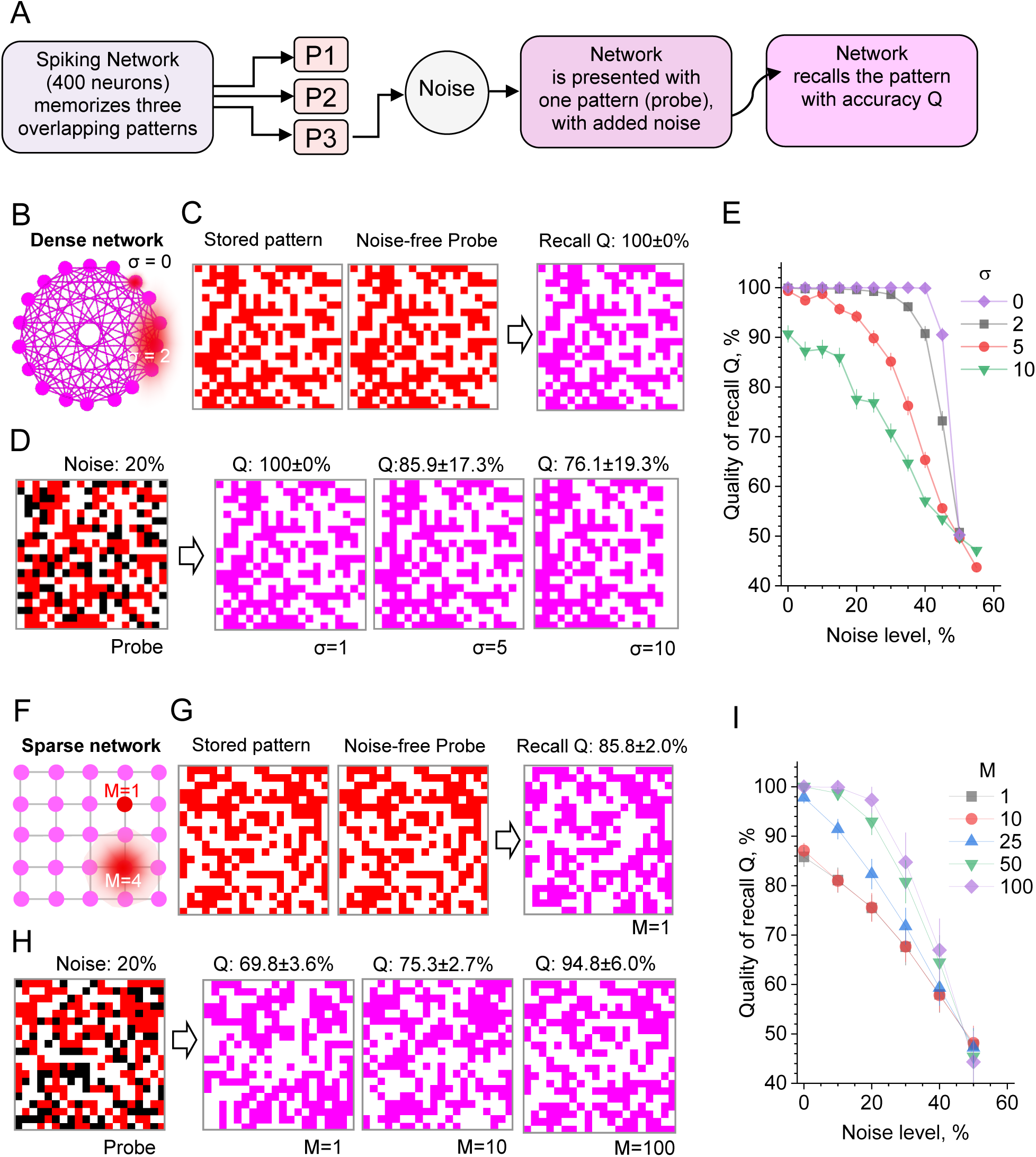
Inter-synaptic glutamate-mediated crosstalk improves memory recall in sparse neural networks. **(A)** Simulation protocol, in which a network is first made to memorize three arbitrary patterns (P1-P3, Figure S10A) and then presented with one of them as a Probe (cue), with random noise added; the network recalls the presented Probe with varied accuracy. **(B)** Schematic illustration of an artificial spiking Generalized Hopfield Network (*N* = 400 cells) with full connectivity; illustration only, not to scale. In baseline conditions, each cell excites all other cells through one-to-one synaptic connections. Glutamate-mediated crosstalk (red shade) is introduced as partial, spatially decaying activation of neighboring cells by synapses activated on the target cell: parameter σ denotes virtual Gaussian dispersion (exponential decay) of excitation, reflecting the number of the affected nearest-neighbor neurons (σ = 0 corresponds to no spillover); see STAR Methods for further detail. **(C)** Example, memory-recall performance of the Hopfield neural network. The network storing three patterns (Figure S10A) is presented with one of them as shown (red pixels, excited neuron states), as a *Probe* pattern, under no spillover (σ = 0); see examples of synaptic weight matrix and spiking raster plot in Figure S10B-C. Unsupervised, the network generated the *Recall* pattern (magenta) which recalls the *Stored* pattern with perfect quality Q = 100% (n = 10 trials; see STAR Methods for definitions). **(D)** Examples of memory recall under noisy *Probe* and excitation spillover. Simulation protocol similar to **C**, but with the *Probe* pattern is contaminated with 20% by noise (black pixels). Under varied excitation spillover (σ = 1, 5, 10), the unsupervised network generated the *Recall* patterns (magenta) reproducing the *Probe* pattern with the quality-of-recall *Q* (mean ± SEM, n = 10 trials for each σ value). **(E)** Summary of simulation experiments shown in **C-D**. Quality of recall *Q* plotted against the noise level in the *Stored* pattern used as a *Probe*, for varied extent of the excitation crosstalk σ. **(F)** Schematic of a spiking neural network with sparse connectivity, in which each cell is connected only to three neighbors; not to scale (cell links do not necessarily indicate reciprocal connections); as in **B,** glutamate-mediated crosstalk (red shade) is introduced as partial, spatially decaying activation of up to M neighboring cells, with the Gaussian dispersion; see STAR Methods and for further detail and Figure S10D-E for examples of synaptic weight matrix and spiking raster plot . **(G)** Example, memory-recall performance in a sparsely connected neural network in noise-free conditions and no crosstalk; other notations as in **C**. **(F)** Examples of memory recall in a sparsely connected network, under noisy input and varied extent of excitation spillover (M = 1, 10, 200); other notations as **D**. **(G)** Summary of simulation experiments shown in **G-H**; other notations as in **E**.

We used classical fully connected (dense) networks ^87,88^ consisting of 400 neurons, with 20 x 20 binary patterns stored as memories (Figure S10A) after a short period of network training (Figure S10B-C). In these networks, glutamate-mediated inter-synaptic crosstalk was represented as an additional, distance-dependent (fading) excitatory input to neighboring cells accompanying each direct excitatory connection (Figure 5B; STAR Methods). The extent of this simulated crosstalk was guided by our experimental model, in which the volume density of excitatory synapses in L1 (∼0.79 μm^−3^) ^55,56^ corresponds to ∼18 and ∼68 neighbors at 1.5 and 2.5 µm, respectively, from individual synaptic connections.

Simulations showed that dense networks showed perfect recall in noise-free conditions, in the absence of inter-synaptic crosstalk (Figure 5C). Introducing the latter indeed appeared to impair their memory, especially in conditions of noisy cues (Figure 5D-E).

However, neural circuits in the brain exhibit very sparse functional connectivity, typically in the low-percent range ^89,90^, with many neuronal groups lacking direct reciprocal connections. To better approximate this architecture, we repeated the memory retrieval tests in sparsely connected networks, in which each cell was connected to only three other cells (Figure 5F, Figure S10D-E). Remarkably, under these conditions, synaptic crosstalk improved memory recall by up to 30%, even for noise-free probes, and remained beneficial until noise contamination exceeded ∼40% (Figure 5G-I). These findings provide proof-of-principle theoretical support for the idea that glutamate spillover can, in certain conditions, enhance recognition capacity in sparsely connected neural networks characteristic of the living brain.

## DISCUSSION

### Principal findings

The present study combines multiplexed optical monitoring of identified thalamocortical synapses to examine how sensory experience reshapes cortical connectivity. We show that rhythmic whisker stimulation induces a robust increase in presynaptic glutamate release efficacy at active POm synapses while recruiting previously silent thalamocortical connections. At the same time, sensory stimulation generates substantial extrasynaptic glutamate transients extending beyond individual synapses and global extracellular GABA signals spanning large regions of layer 1.

Although RWS-induced LTP had little detectable effect on the overall inhibitory signal, it enhanced the overlap among glutamate transients generated by neighboring excitatory synapses, possibly extending beyond POm projections. Neural network simulations guided by these experimental observations suggest that such glutamate-mediated inter-synaptic crosstalk can improve associative memory retrieval in sparsely connected networks.

Together, these findings indicate that activity-dependent plasticity may modify not only the efficacy of anatomically defined synaptic pathways but also the extent of functional communication among nearby synapses, thereby dynamically reshaping the operational synaptic connectome.

### Activity-driven synaptic plasticity in vivo

It is widely accepted that information transfer and storage in the brain are encoded largely through changes in synaptic weights that follow Hebbian learning principles. Recent advances in targeted molecular labelling of synaptic activity have revealed memory-associated changes in the brain connectome, involving alterations in circuit efficacy, synaptic number, and connectivity patterns ^17,18,91^. While mapping accumulated activity across defined synaptic populations has provided important insights into the concept of the memory engram, monitoring the efficacy of identified individual synapses, and its experience-dependent modulation in vivo, remains technically challenging. Yet, distinguishing the relative contributions of postsynaptic current enhancement and changes in release probability (P_r_) is essential for understanding the cellular mechanisms through which experience reshapes the functional connectome.

In this context, physiological LTP paradigms in vivo remain an important tool for dissecting the mechanisms of synaptic plasticity. Although enhancement of postsynaptic responsiveness is often regarded as the canonical expression mechanism of LTP, presynaptic plasticity, both short-and long-term, appears equally widespread throughout the mammalian brain ^30,92–94^. This is perhaps unsurprising because the 50-150% increase in postsynaptic current typically associated with LTP can, at least in principle, be exceeded by substantially larger changes in P_r_.

While accurate measurements of P_r_ can be achieved in controlled conditions using optical glutamate sensors in cultured neurons or brain slices ^25,58,59^, responses to physiological stimulation in vivo are inherently noisy and difficult to relate directly to presynaptic spike output. We therefore introduced a ratiometric measure of synaptic release efficacy, E_r_, defined as the ratio between evoked glutamate and presynaptic Ca²⁺ signals. Control experiments indicated that, under non-saturating sensor conditions, this metric scales with the amount of glutamate released per unit presynaptic activity. Using this approach, we found that RWS-induced LTP increased E_r_, reflecting a combination of enhanced glutamate release and reduced presynaptic Ca²⁺ responses to brief sensory stimulation. These observations support a significant presynaptic component of thalamocortical plasticity, manifested as increased glutamate output per unit presynaptic activity.

At the same time, the increase in glutamate release at individual synapses was insufficient to account fully for the magnitude of potentiation detected electrophysiologically. This prompted us to examine the population of monitored POm boutons, revealing that RWS-LTP also recruited a substantial proportion of previously silent thalamocortical connections. The fraction of RWS-responsive boutons increased by nearly 50%, indicating that sensory experience can rapidly reconfigure the distribution of active inputs within an existing anatomical projection. These findings suggest that physiological plasticity in vivo is not restricted to modifying the strength of already active synapses but may also involve dynamic allocation of functional connectivity through the engagement of previously ineffective pathways. In this respect, the present observations complement emerging evidence that memory-associated plasticity entails not only changes in synaptic efficacy but also reorganization of the active synaptic ensemble supporting information processing and storage.

### Detecting glutamate spillover in the intact brain

Glutamate escaping the synaptic cleft is rapidly bound by high-affinity transporters ^31,32^, predominantly GLT-1 expressed by astrocytes at an estimated surface density of 1-2·10^4^ µm^-2^ (refs. ^33,95^). This implies the presence of 1-2·10^5^ GLT-1 molecules within one micron of a release site, dwarfing the estimated 2,000-8,000 glutamate molecules discharged by a single synaptic vesicle. Accordingly, biophysical models incorporating the overall buffering capacity of GLT-1 in perisynaptic neuropil have predicted minimal glutamate diffusion beyond ∼0.5 µm from the release site ^38,76,96–99^.

However, multiple electrophysiological studies in brain slices suggested that glutamate can activate receptors beyond one micron from it release site, a phenomenon subsequently corroborated using genetically encoded glutamate sensors ^25,40^. A likely explanation for the discrepancy between these observations and earlier theoretical predictions is that most models distributed glutamate transporters uniformly throughout the neuropil at their average density, whereas astroglial processes occupy only 8-10% of neuropil volume and closely approach only ∼60% of hippocampal synapses ^100,101^. In contrast, a model in which GLT-1 is concentrated on astrocyte membranes, while maintaining the same overall transporter density, readily predicts glutamate escape beyond one micron (Figure 4G-I).

Importantly, the present observations extend this concept to the intact brain. High-resolution imaging of astrocyte-expressed iGluSnFR revealed glutamate transients extending over distances comparable to, or exceeding, the average spacing between neighboring excitatory synapses in layer 1. Under these conditions, the measured signal is unlikely to reflect glutamate released by a single synapse in isolation. Instead, it is more likely to represent the integrated footprint of multiple nearby glutamatergic events, including activity arising from neighboring thalamocortical and intracortical connections.

This interpretation is supported by two observations. First, restricting iGluSnFR expression to individual POm boutons revealed a markedly more confined glutamate profile, closely matching previous measurements obtained in brain slices. Second, RWS-induced LTP increased the long-range component of the extrasynaptic glutamate signal without measurably altering the local escape profile surrounding individual potentiated synapses. Together, these findings suggest that sensory plasticity enhances the overlap among glutamate transients generated by neighboring synapses rather than increasing glutamate escape from individual release sites.

Such enhanced overlap is consistent with both the recruitment of previously silent thalamocortical inputs and the withdrawal of perisynaptic astroglial processes observed after RWS-LTP, two mechanisms that would be expected to increase the probability of inter-synaptic glutamate interactions. More broadly, these observations suggest that activity-dependent plasticity can alter not only the efficacy of individual synapses but also the extent to which neighboring synapses communicate through the extracellular space, thereby reshaping the operational connectivity of local cortical circuits.

### Functional connectivity beyond anatomical wiring

The classical concept of a neural connectome is rooted in anatomically defined synaptic connections. Within this framework, information is transmitted through discrete point-to-point contacts whose efficacy can be modified by activity-dependent plasticity. The present findings suggest that this picture may be incomplete. In addition to altering the strength and recruitment of synaptic connections, sensory experience appears capable of modifying the extent of glutamate-mediated communication among neighboring synapses that are not necessarily connected anatomically.

This distinction is important because the extracellular space constitutes a shared signaling environment for large numbers of nearby synapses. A brief RWS appears to trigger glutamate release from thalamocortical synapses that could exert influence beyond their immediate vicinity. Following RWS-induced plasticity, the combination of enhanced release efficacy, recruitment of previously silent inputs, and withdrawal of perisynaptic astroglial processes is predicted to increase the overlap among neighboring glutamate transients. As a consequence, receptors located outside the activated synaptic pathway may become exposed to greater glutamate concentrations sufficient for ligand binding and, potentially, receptor activation.

These observations support a view in which the effective communication landscape of a neural circuit is shaped not only by its anatomical wiring diagram but also by the spatiotemporal dynamics of extracellular signaling. In this framework, synaptic plasticity modifies both the weights of existing connections and the probability of interaction among neighboring synapses through volume-transmitted signals. Such a mechanism would provide a dynamic means of expanding or contracting functional connectivity without requiring structural rewiring of the underlying circuit.

### Glutamate spillover and network memory performance

The computational consequences of glutamate spillover remain poorly understood. From a traditional perspective, inter-synaptic crosstalk might be expected to degrade information transfer by blurring the boundaries between anatomically distinct connections. Indeed, this intuition is consistent with our simulations of densely connected Hopfield networks, in which volume-transmitted excitation impaired memory retrieval, particularly under noisy conditions. In such networks, individual neurons already receive extensive direct connectivity, leaving little scope for additional excitatory interactions to improve information transfer.

The situation was markedly different in sparsely connected networks, which more closely resemble the low connection probabilities characteristic of cortical circuits ^89,90^. Under these conditions, introducing glutamate-mediated crosstalk improved associative memory retrieval across a broad range of parameters, including noise-free inputs. Although highly simplified, these simulations suggest that local volume transmission can compensate, at least in part, for sparse anatomical connectivity by providing additional routes for information flow between neuronal ensembles.

It is important to emphasize that the present simulations were not intended to reproduce the architecture of barrel cortex or to provide a realistic model of memory storage in vivo. Rather, they were designed to test a general computational principle emerging from our experimental observations: whether excitatory communication extending beyond anatomically connected pathways could influence network performance. Within this framework, glutamate spillover acts not as a source of noise but as a mechanism that transiently increases effective connectivity among neighboring network elements.

Viewed in this way, the recruitment of previously silent thalamocortical inputs and the increased overlap among glutamate transients observed after RWS-induced plasticity may represent complementary aspects of the same process. Both mechanisms effectively expand the pool of interactions available to the network without requiring structural modification of the underlying circuitry. While the extent to which cortical circuits exploit such mechanisms remains unknown, the present findings raise the possibility that extrasynaptic signaling contributes to information processing in ways that are not readily captured by conventional connectomic descriptions of neural networks.

### Limitations

Several limitations should be considered when interpreting the present findings. First, glutamate sensors act as additional glutamate-binding sites and therefore inevitably perturb the extracellular environment they are used to measure. Although our simulations explicitly incorporated the buffering effects of iGluSnFR and indicated that the resulting estimates of glutamate escape are conservative, the true spatiotemporal profile of extracellular glutamate in unperturbed tissue remains difficult to establish experimentally. Likewise, while the agreement between bouton-targeted iGluSnFR measurements and previous observations in brain slices provides confidence in our estimates, the spatial distribution and expression level of the sensor could vary between preparations and across cellular compartments.

Second, the present experiments were conducted in a specific thalamocortical pathway and employed a well-established paradigm of sensory plasticity. The extent to which the observed relationship between synaptic potentiation, astroglial remodeling, glutamate spillover, and recruitment of previously silent inputs generalizes to other cortical circuits, developmental stages, or behavioral contexts remains unknown. Similarly, although our analyses suggest that extrasynaptic glutamate can reach concentrations sufficient for ligand binding and potentially receptor activation at neighboring synapses, direct measurements of receptor activation outside the stimulated pathway were beyond the scope of the present study.

Finally, the network simulations were intended to explore a computational principle rather than to reproduce the architecture of barrel cortex or the mechanisms of memory storage in vivo. The implementation of glutamate-mediated crosstalk as a distance-dependent increase in effective connectivity represents a simplified abstraction of a complex biological process involving neuronal geometry, receptor distribution, transporter dynamics, and ongoing network activity. The observed improvements in memory retrieval should therefore be interpreted as proof-of-principle rather than evidence that cortical circuits exploit glutamate spillover specifically for memory storage or recall.

Despite these limitations, the convergence of independent experimental observations and theoretical analyses supports the conclusion that activity-dependent plasticity can influence not only the strength of anatomically defined synaptic pathways but also the extent of communication among neighboring synapses through the extracellular space.

### Concluding remarks

The present findings suggest that experience-dependent plasticity can modify cortical information processing through multiple complementary mechanisms. In addition to increasing glutamate release efficacy at active synapses and recruiting previously silent inputs, sensory stimulation enhanced the overlap among glutamate transients generated by neighboring excitatory connections. These observations support the idea that synaptic plasticity may alter not only the strength of anatomically defined pathways but also the extent of communication among nearby synapses through the extracellular space. Such activity-dependent changes could dynamically reshape the effective interaction landscape of local neural circuits without requiring structural rewiring of the underlying connectome. More broadly, these findings suggest that the operational connectivity of neural circuits may be determined not only by anatomical synaptic wiring but also by dynamic interactions mediated through the extracellular environment.

## RESOURCE AVAILABILITY

### Lead contact

Requests for further information and resources should be directed to and will be fulfilled by the lead contact, Dmitri Rusakov.

### Materials availability

This study did not generate new unique reagents.

### Data and code availability

Original data from electrophysiology and imaging experiments are fully available upon request. The computer modelling results presented here could be directly replicated using the simulation software, including programming code, deposited at https://github.com/RusakovLab/DiffusionPlusNeuralNetwork, with the relevant study parameters outlined in the main text and figures.

## Acknowledgments

This work was supported by reserach grants from Wellcome (101896/Z/13/Z, 223131/Z/21/Z), Medical Research Council (MR/W019752/1), NC3Rs (NC/X001067/1), and BBSRC (BB/Y003926/1, BB/Y009959/1) to D.A.R; and UCL-Wellcome Pilot Award (RM-TIN OK: #178973) to O.K.

## Author contributions

O.K. and J.P.R. set up and carried out in vivo experiments; T.P.J. carried out experiments in brain slices and data analyses; O.K., J.P.R., and T.P.J. carried out viral transduction; K.Z. anlaysed in vivo imaging data; L.P.S. designed and carried out biophysical and neural-network simulations; D.A.R. conceptualized and narrated the study, carried out data analyses, compiled illustraitons, and wrote the original manuscript, which was further contributed to by all authors.

## STAR METHODS

### EXPERIMENTAL MODEL AND STUDY PARTICIPANT DETAILS

#### Animal experimentation

All animal procedures were conducted in accordance with the European Commission Directive (86/609/ EEC), the United Kingdom Home Office (Scientific Procedures) Act (1986) with project approval from the Institutional Animal Care and Use Committees of the University College London. All animals were maintained in controlled environments as mandated by national guidelines, on 12hr light/dark cycles, with food and water provided ab libitum.

#### Experimental model designs across preparations

For ex vivo electrophysiology and imaging both male and female C57BL/6 J mice (Charles River Laboratories) were used. For experiments requiring viral-mediated expression male and female wild-type C57BL/6 mice (Charles River Laboratories) were injected at P0-1 days of age with viral vectors, and acute brain slices were obtained on average four weeks later. For experiments *in vivo*, male and female wildtype C57BL/6 mice (Charles River Laboratories) were injected with viral constructs at 1-1.5 month old; all animals underwent craniotomy and the implantation of a head plate at 4 to 8-weeks post-injection, as detailed below.

### METHOD DETAILS

#### Organotypic Slice Preparation

Organotypic hippocampal slice cultures were prepared and grown with modifications to the interface culture method ^102^ from P6–8 Sprague-Dawley rats, in accordance with the European Commission Directive (86/609/EEC) and the United Kingdom Home Office (Scientific Procedures) Act (1986). Three hundred μm thick, isolated hippocampal brain slices were sectioned using a Leica VT1200S vibrotome in ice-cold sterile slicing solution consisting (in mM) of Sucrose 105, NaCl 50, KCl 2.5, NaH2PO4 1.25, MgCl2 7, CaCl2 0.5, Ascorbic acid 1.3, Sodium pyruvate 3, NaHCO3 26 and Glucose 10. Following washes in culture media consisting of 50% Minimal Essential Media, 25% Horse Serum, 25% Hanks Balanced Salt solution, 0.5% L-Glutamine, 28mM Glucose and the antibiotics penicillin (100U/ml) and streptomycin (100μg/ml), three to four slices were transferred onto each 0.4μm pore membrane insert (Millicell-CM, Millipore, UK), kept at 37°C in 5% CO2 and fed by medium exchange for a maximum of 21 days in vitro (DIV).

#### Biolistic Transfection

The second generation iGluSnFR variant SF-iGluSnFR.A184V, kindly gifted by Prof. Loren Looger was expressed in CA3 pyramidal cells in organotypic slice cultures using biolistic transfection techniques adapted from manafacturer’s instructions. In brief, 6.258 mg of 1.6 μm Gold microcarriers were coated with 30 µg of hSyn-SF-iGluSnFR.A184V and or hSyn-axon-jRGECO1a plasmid. Organotypic slice cultures at 5DIV-6DIV were treated with culture media containing 5µM Ara-C overnight to reduce glial reaction following transfection. The next day cultures were shot using the Helios gene-gun system (Bio-Rad) at 120psi. The slices were then returned to standard culture media the next day and remained for 5-10 days before experiments were carried out.

#### Axon tracing and 2PE imaging in axonal boutons

We used a Femtonics Femto2D-FLIM or a Femto3D-RC-FLIM imaging system, integrated with patch-clamp electrophysiology (Femtonics, Budapest) and linked on the same light path to two femtosecond pulse lasers MaiTai (SpectraPhysics-Newport) with independent shutter and intensity control. Patch pipettes were prepared with thin-walled borosilicate glass capillaries (GC150-TF, Harvard apparatus) with open tip resistances 2.5-3.5 MOhm. For CA3 pyramidal cells and dentate granule cells, the internal solution contained (in mM) 135 potassium methanesulfonate, 10 HEPES, 10 di-Tris-Phosphocreatine, 4 MgCl_2_, 4 Na_2_-ATP, 0.4 Na-GTP (pH adjusted to 7.2 using KOH, osmolarity 290–295), and supplemented with Cal-590 (300 µM; AAT Bioquest) for FLIM imaging in CA3 pyramidal cell axons.

Presynaptic imaging at CA3-CA1 synapses was carried out using an adaptation of presynaptic glutamate and Ca^2+^ imaging methods previously described ^25^. CA3 pyramidal cells were first identified as iGluSnFR expressing using 2PE imaging at 910 nm and patched in whole cell mode as above. Following break-in, 30-45 minutes were allowed for Cal-590 or Alexa 488 (200 µM) to equilibrate across the axonal arbor (Figure 1I). Axons, identified by their smooth morphology and often torturous trajectory were followed in frame scan mode to their targets and discrete boutons were identified by criteria previously demonstrated to reliably match synaptophysin labelled punctae ^103^. In some cases, 4µM Alexa 594 was included with Cal-590 in the internal solution (Figure S3). The distinct two-photon excitation profiles of these two red emitting dyes enable morphology to be traced by Alexa 594 emission with 800nm excitation, and Ca^2+^ signals to be recorded in the same structure by Cal-590 emission at 910nm excitation, with no significant contribution of Alexa fluorescence to the Cal-590 emission.

For fast imaging of action-potential evoked iGluSnFR, jRGECO1a, or Cal-590 fluorescence transients at single boutons, a spiral shaped (“Tornado”) scan line was placed over the bouton of interest (described further in the text) which was then scanned at a sampling frequency of ∼500 Hz with excitation at 910 nm. For multi-bouton imaging point-scans were made with a temporal resolution ∼333 or 250 Hz, usage described further in the text. Following a baseline period, action potentials initiated by brief positive voltage steps in voltage clamp mode (V_m_ holding -70 mV) were given with an interval of 50 ms.

#### Viral transduction of jRGECO in POm neurons and iGluSnFR in cortical astrocytes

All animal procedures were conducted in accordance with the European Commission Directive (86/609/EEC) and the United Kingdom Home Office (Scientific Procedures) Act (1986). Adult C57BL/6J mice, male and female, were transfected with two viral constructs, encoding hSyn.jRGECO1a and GFAP.SF-iGluSnFR.A184S respectively. Mice were anaesthetised (isoflurane, maintenance at 1.5 - 2%), prepared for aseptic surgery and secured in a stereotaxic frame. Upon confirmation of the absence of pedal withdrawal reflex, two craniotomies of approximately 0.4 mm diameter were performed over the right hemisphere using a high-speed hand drill (Proxxon, Föhren, Germany), at sites overlying the posterior medial nucleus of the thalamus (POm) and the barrel cortex (S1BF). The entire microinjection into the POm was completed prior to performing the second craniotomy over S1BF. Stereotactic coordinates for POm injections (jRECO1a) were -2.2 mm and 1.2 mm along the anteroposterior and mediolateral axes, respectively. Two injection boluses were delivered at 2.9 and 3.1 mm beneath the dural surface. For S1BF injections (iGluSnFR), the bolus coordinates were -0.5 mm and 3.0 mm along the anteroposterior and mediolateral axes, respectively, delivered at a depth of 0.6 mm along an oblique approach angle to avoid tissue scarring in S1BF that might occlude the optical window. A warmed saline solution was applied to exposed cortical surface during the procedure.

Pressure injections of AAV9 hSyn.jRGECO1a (totalling 0.5 x 10^10^ genomic copies in a volume not exceeding 200 nL, initially supplied by Penn Vector Core, PA, USA; and later by Addgene, MA, USA) and AAV2/5.GFAP.iGluSnFR.A184S (0.1 x 10^10^ genomic copies, in a volume not exceeding 200 nL, kindly gifted by Prof. Loren Looger) were carried out using a glass micropipette at a rate of 1 nL sec-1, stereotactically guided to the POm and S1BF, respectively, as outlined above. Once delivery was completed, pipettes were left in place for 5 minutes before being retracted. The surgical wound was closed and the animal recovered in a heated chamber. Meloxicam (subcutaneous, 1 mg kg^-1^) was administered once daily for up to two days following surgery. Mice were subsequently prepared for cranial window implantation approximately 2-3 weeks later.

#### Dual transduction of jRGECO and iGluSnFR in POm neurons

Animals received one injection of two viral constructs, encoding hSyn.jRGECO1a and hSyn.iGluSnFR.A184V (Addgene, MA, USA), given using a Hamilton syringe stereotactically guided to target the POm following the coordinates as specified above. The injection was made at a depth of 3 mm beneath the cortical surface, delivered a single bolus under control of a microinjection pump, in a volume not exceeding 200 nL. Perioperative multimodal analgesia was carried out with buprenorphine (60 μg kg^-1^, s.c.) and lidocaine (2.5%) topically applied to the surgical site; metacam (1 mg kg^−1^, s.c.) and ocular ointment (Lacri-lube, Allergan, UK) were also applied. Isoflurane was used for anesthesia throughout the surgical procedure: 4% v/v for induction and 1.5-2.5% v/v for maintenance. Body temperature was maintained at ∼37.0 °C using a feedback rectal thermometer and heating blanket. After the surgical wound was closed, saline (0.5 mL, s.c.) was administered, and the animal was left to recover in a heated chamber. Animals were monitored for several days perioperatively or until the wound healed.

#### Headplate installation, craniotomy, durotomy

Animals were prepared for craniotomy as described for the viral injection procedure 3–4 weeks after dual transduction of viral vectors. After an animal was secured and deeply anesthetized, the skull’s right frontal and parietal bones were exposed; the area was cleaned and coated with tissue adhesive (3M Vetbond, UK) to facilitate headplate installation. A custom-made headplate was affixed over the right somatosensory cortex (the targeted injection site) and secured with dental cement (SuperBond, Sun Medical Co. Ltd., Japan). Once the cement components had cured, the animal was secured in a custom-built head fixation frame. A craniotomy of ∼3 mm diameter was performed over the S1BF region using a high-speed hand drill. After sufficiently thinning the skull and superfusing its surface with saline, the skull flap was removed using fine-tipped forceps. Immediately after opening, the brain was superfused with sterile saline. Durotomy was carried out using 28G needles with hand-made curved tips, avoiding penetrating or damaging the pia mater. After completing the craniotomy, the cranial window was securely closed with a double-glass – custom-prepared assembly consisting of two round coverslips of a 3 mm and 4 mm diameters (Harvard Apparatus UK) glued together – and affixed over the exposed cortical region as previously described ^104^. Slight downward pressure was applied to the coverslip assembly using a stereotactically guided wooden spatula to allow some flexible force while securing the glass window with dental cement. Once the cement had cured, the animal was transferred to the microscope with a heated blanket and controlled temperature for multiplex imaging.

In some experiments, the anaesthesia regime was switched from isoflurane to a mixture of fentanyl (0.03 mg kg-1, i.p.), midazolam (3 mg kg-1), and medetomidine (0.3 mg kg-1), for the subsequent high-resolution multiplexed two-photon imaging in the anaesthetised animal.

#### Electrophysiological control of RWS-LTP induction

For electrophysiological recordings, the cranial window was not closed with the glass window. Extracellular local field potential (LFP) recordings in barrel cortex were obtained using standard glass micropipettes (Warner Inst.) of low resistance (1.5–2.5 MΩ) filled with an extracellular Ringer solution. The recording pipette was positioned at the level of L1-2, typically at approximately -100 µm depth under visual control to prevent any damage to blood vessels, avoiding lateral movement of the pipette once within the tissue. For somatosensory stimulation, the contralateral whiskers were stimulated with pressurised nitrogen to evoke tactile responses (rhythmic whisker stimulation, RWS) within the regions of interest. The baseline stimulus was set by probing individual whisker stimulation to ensure stable responses to a burst of RWS (4 pulses applied at 20 Hz); control RWS trains were applied every 30 sec, for at least 15 min to ensure stable responses. RWS-evoked LTP was induced using two trains of low-frequency continuous whisker stimulation (3 Hz for 60 s each). The control RWS responses were monitored afterwards for at least an hour using the same protocol (4 pulses at 20 Hz). Recordings were carried out using a Multiclamp 700B controlled by pClamp 10.2 software (Molecular Devices). Signals were filtered at 3-10 kHz, digitized and sampled using a Digidata (Molecular Devices) at 10-20 kHz. The responses were analyzed offline using Clampfit 10.3 software (Molecular Devices) as area under the curve (AUC) relative to baseline responses.

#### Multiplexed 2PE imaging in vivo

Multiplexed two-photon excitation imaging was carried out in two settings. The first setting comprised a FemtoSmart imaging system (Femtonics, Budapest) integrated with electrophysiology and incorporating a SpectraPHysics Insight X3 dual-output laser, combined galvo-galvo and resonant scanner scanhead as detailed previously ^105^. The anaesthetised animal was secured on a custom-built stage via the installed headplate under XLPlan *N* 25× water immersion objective (NA 1.05) coupled to a green lamp illumination. The second setting used an Olympus FV1000 microscope, a wavelength multiplexing suite integrated with electrophysiology, which consisted of a Newport-Spectraphysics Ti:sapphire MaiTai tunable IR laser pulsing at 80 MHz and a Newport-Spectraphysics HighQ-2 fixed-wavelength IR laser pulsing at 63 MHz, as detailed earlier ^48,106,107^. The laser light paths were aligned (though not synchronized) before being point-scanned using an XLPlan N 25x water immersion objective (NA 1.05). Imaging was performed with head-fixed, awake animals as well as under a lightly anesthesia (low doses of isoflurane) or a deeply anesthetized regimen (fentanyl, 0.03 mg kg^-1^, midazolam, 3 mg kg^-1^, and medetomidine, 0.3 mg kg^-1^).

First, exploratory acquisitions were performed with both lasers illuminating the tissue at 920 nm and 1045 nm, respectively, in order to locate thalamocortical axons in S1BF ramifying within the arbor of iGluSnFR-positive cortical astrocytes. The contralateral whiskers were stimulated (RWS; 5 seconds, 3 Hz) to confirm tactile responses within the regions of interest. Measurements were performed throughout in L1 and sometimes L2, at depths of 50-200 μm. For high resolution axonal bouton sampling, an initial frame-scan of 4-20 Hz was performed, with a pixel dwell time of 2 μs and a mean laser power not greater than 30 mW at the focal plane. At this stage, responses to short sensory stimulus trains (20 Hz, 200-ms) were monitored at axonal boutons, both before and after RWS-induced LTP while in Tornado scanning mode (400-500 Hz). RWS-evoked LTP within the barrel cortex was induced as described above and previously ^39,44^, via a sustained contralateral RWS (120 sec, 3 Hz).

Overall, our sample included 44 individual synapses in LTP experiments across 16 animals. In an additional 10 animals, LTP experiments could not be carried out due to insufficient signal in one or both imaging channels.

#### Analyses of optical recordings: frame scan, Tornado scan, resonant scan

Optical recordings were acquired using one of three imaging modes: (i) frame time series with a galvo-mirror scanhead (10–30 Hz), (ii) Tornado spiral linescan (0.4–1 kHz), or (iii) resonant-mirror frame scanning (0.2–0.4 kHz; Femtonics FemtoSmart system only).

Galvo-mirror frame scanning was used to identify fields of view under optimal conditions for multiplexed dual-channel imaging and detection of responsive synaptic connections (Figure S1). Owing to its relatively low temporal resolution and signal noise, only eight synaptic connections met the criteria for subsequent analysis.

Tornado linescan recordings were analyzed using two complementary approaches: one focused on temporal signal dynamics, and the other on spatial signal features. In the first approach, fluorescence signals were integrated across each individual scan (0.5-1 kHz) and stored as time-course traces (Figure 2f,h). Prior to analysis, photobleaching and other baseline trends, when present, were corrected by subtracting an exponential or linear fit applied to the trace before and after the RWS-induced response. XY-axis motion artefacts were corrected in the recorded image series; any data that cannot be reliably corrected were excluded from the analysis. All analyses were performed on unfiltered data. For illustrative clarity, two types of unsupervised signal filtering were subsequently applied: (i) a low-pass filter to suppress optical noise caused by rapid brightness fluctuations from stochastic photon emission during the microsecond pixel dwell time, and (ii) a band-pass filter to reduce brightness fluctuations associated with the animal’s heartbeat during seconds-long recordings.

In the second approach, Tornado linescan datasets were reconstructed into near-circular two-dimensional fluorescence landscapes representing the scanned spiral area (Figure 3d-f, Figure S7) (available at https://github.com/tomjensen2/Fiji-GUI) . Because the lateral distance between adjacent spiral turns was below the diffraction limit (∼0.2 µm), corresponding pixels were concatenated without appreciable loss of optical resolution. Time series of reconstructed images were used to obtain averaged fluorescence landscapes over defined time windows, such as 200-ms before and 200-ms during the RWS-induced response (Figure 4A-B, D; Figure S7). Analyses were performed on the original unfiltered images.

Resonant-scanner frame imaging was used to record high-speed fluorescence dynamics from multiple axonal boutons (Figure S2E-F) or across entire regions of interest (Figure 3H-I). The microsecond pixel dwell time in these recordings produced substantial frame-to-frame emission fluctuations within small ROIs. Consequently, 0.2-0.3 kHz time series were binned (averaged) five-fold for RWS-induced trials and two-fold for spontaneous activity detection. For LTP measurements, fluorescence responses to 200-ms RWS stimuli were averaged over 5-10 trials collected 15-1 min before induction and 10-30 min after induction, in both imaging channels.

#### Analyses of spontaneous events

Spontaneous synaptic events were recorded in a subset of targeted connections in which the signal-to-noise ratio enabled reliable detection, normally in either iGluSnFR or jRGECO1a channel. Detection and analysis of events were carried out using either Mini Analysis 6 (Synaptosoft) or open-source Easy Electrophysiology v2.7.3 (https://www.easyelectrophysiology.com/), as detailed in Figure S2a-d. Spontaneous events analyzed in figure 3G-H were initially manually selected through visual inspection of the spiral scanning sequence temporal trace. Release hotspot was then localised at each time point as previously described ^25^. Localization hotspot at the edge of or outside the bouton profile, with slow rising time, possibly due to spillover from neighboring synapses, were rejected.

#### Simulations of glutamate escape: synaptic microenvironment

The MATLAB-based modelling framework was previously developed, validated, and constrained across multiple experimental paradigms ^75,78,108,109^. The simulated arena consisted of a 4-μm-wide cube containing a central synaptic structure represented by a 220-nm-wide, 20-nm-high synaptic cleft separating two hemispherical, pre-and postsynaptic, diffusion obstacles ^77,80^ (Figure S9a). Surrounding this synaptic configuration were stochastically generated structures representing neuronal and astrocytic processes, occupying approximately 70% and 10% of the tissue volume, respectively, in barrel cortex layer 1 ^73,74^, leaving ∼20% for the extracellular space ^81,82^. These cellular processes were modelled as randomly sized, overlapping, and concatenated spheres, often forming caterpillar-like chains, as described previously ^75,110^. This approach, firstly, enables each simulation run to produce an ‘accidental’ perisynaptic neuropil configuration, reflecting the inherent variability of individual synapses. Secondly, it generates a diverse mixture of concave and convex geometries, including diffusion ‘dead-ends,’ an established structural feature of brain neuropil ^111^.

Space filling with overlapping spheres followed the procedure described previously ^109^. In brief, the key control parameter was the volume fraction (β) occupied by the spheres defined as β = 1−α, where α stands for the extracellular space fraction (α = 0.2 in our case). To generate a distributed set of overlapping spheres, random 3D coordinates for sphere centroids were assigned throughout the arena, along with a random radius for each sphere. Sphere radii followed a uniform distribution between 50 and 300 nm, approximating the characteristic widths of cortical neuropil elements (fragments of neuronal and astroglial processes) observed in electron micrographs. For each simulation run, the initial number of spheres was estimated based on their average size and volume to yield an approximate total occupancy of β = 0.8. The central synaptic structure was excluded from sphere placement. The value of β was estimated by (a) uniformly scattering 10^5^ test points throughout the simulation arena and (b) calculating the proportion of points falling outside the spheres. Increasing the number of test points to 10^6^ changed β by less than 1%, indicating asymptotic accuracy. The space-filling procedure was then repeated, with the number of spheres adjusted iteratively, until β converged to the target value within approximately 5% accuracy.

#### Simulations of glutamate escape: diffusion and binding

In each simulation run, up to 5000 glutamate particles were released instantaneously at the synaptic cleft center. Inside and outside the cleft, particles diffused through the tortuous extracellular space with a free-space diffusivity of *D* = 0.5 μm^2^/ms, as previously determined using anisotropy-FLIM ^106^. The actual value of *D* was regularly verified to prevent artefacts arising from computational “traps” in Monte Carlo algorithms.

Interactions between individual particles and spheroidal structures representing either neuronal or astroglial fragments were modelled in two distinct ways. For ’neuronal’ spheroids, collisions were treated as elastic reflection. For the ’astroglial’ spheroids representing cell surfaces density populated with high-affinity GLT-1 transporters and/or iGuSnFR, interactions were modelled as binding events occurring with a probability *P_b_*. For each particle, *P_b_* was a function of the elapsed time (*t*) since its first encounter with the present astroglial surface, following the lifetime expression for a first-order reaction: *P_b_*=1−exp(−*t ·*Ψ^−1^), where Ψ is the time constant governing the likelihood of binding. Parameter Ψ effectively combines binding affinity and surface density of binding sites, thus representing the astroglial surface binding capacity in the model.

In practice, *P_b_* was computed for each particle as long as it remained within 5 nm of an astroglial spheroid and reset to zero once the particle moved beyond that distance. Increasing the cut-off distance above 5 nm had a negligible effect on *P_b_*. Once bound, particles remained attached to astroglial spheroids for the duration of the simulated time window (4–6 ms), consistent with the much longer characteristic unbinding times (tens of milliseconds) for both GLT-1 and iGluSnFR. The simulation time step, Δ*t* (typically < 0.1 µs), was sufficiently small to prevent particles from “tunnelling” through the smallest 50 nm-wide obstacles.

#### Reconstructing radial 3D distribution from its 2D projection using Abel inversion

The radial distributions of the *ΔF/F_0_* fluorescent emission (Figure 3f-h) represent 2D projections of the signal distribution in 3D. As they have been computed under the assumption of spherical symmetry, the respective 3D distributions can be calculated using the inverse Abel transform:

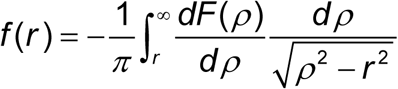

where *F*(ρ) is the radial distribution of the 2D-projected emission signal whose 3D distribution is *f*(*r*). Experimental curves were smoothed with a three-point spline for the transform to converge. The resulting *f*(*r*) distributions (Figure 4c) were normalized for comparison purposes.

#### Simulating Hopfield networks with associative learning and memory retrieval

We explored associative memory retrieval in a recurrent Hopfield network of *N* = 400 binary neurons incorporating a leaky integrate-and-fire model ^112,113^. Each neuron takes one of two activity states per update: active (+1) or inactive (−1). In each trial, the network was trained to store *P_s_* = 3 distinct bipolar patterns encoded as 400-long vectors *ξ ^μ^* ∈ {−1,+1}*^N^* (*µ* = 1, 2, 3), presented as 20 x 20 pixels black-and-white images. The patterns had roughly 40% active (+1) neurons, to keep the memory load well below the classical Hopfield capacity of 0.14*N*, ensuring robust memory retrieval.

The networks were trained to store the patterns using a standard Hebbian rule (co–activation strengthens connections), producing a symmetric synaptic weight matrix **W** with zero self–connections:

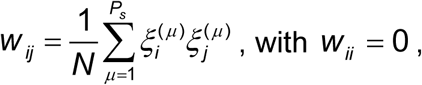

No dilution or sparsification of **W** was applied.

For each trial, the target stored pattern*ξ ^μ^* was randomly selected from the three stored and a noisy probe (cue) *s*_0_ was generated by flipping a fraction *η* (noise level in %) of its entries (*η* ∈ [0, 1]). Recall was initiated from this noisy cue and proceeded through synchronous network updates until convergence.

During memory retrieval process, all neurons were updated synchronously using a standard sign rule ^112,114^. To speed up the process, a weak cue clamp (gain 0.08) was applied during first four iterations, nudging the network towards the target basin ^115^. Up to 200 iterations were run or until the network state was unchanged over five consecutive steps.

For each selected noise level *η*, 100 trials were run each involving three newly generated stored patterns from which one was selected as a noisy probe. The recall quality Q was evaluated by calculating the ratio

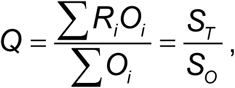

where *O_i_* (i = 1,…,*N*) ∈ {0,1} is the original stored pattern (1 and 0 are targeted active and inactive neuron states) and *R_i_*(i = 1,…,*N*) ∈ {0,1} is the corresponding retrieved pattern. The resulting *S_T_* and *S_O_* values are, respectively, the sum of ’true-positive’ values (neurons active in both stored and retrieved patterns) and of original positive values (neurons active in the original stored pattern).

#### Hopfield networks: introducing synaptic crosstalk (excitation spillover)

Synaptic crosstalk was modelled as the excitation spreading from individual neurons to their neighbors. To implement this, we arranged the network as a virtual ring (Figure S10a illustration) so that the excitation ’spillover’ could be restricted to M neighbours for each neuron. The spillover intensity kernel *k* was shaped by parameter σ, Gaussian dispersion (virtual spatial decay) of the excitation signal, so that

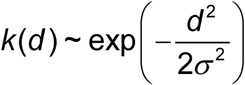

where *d* is the neighbour-neuron distance {1, 2, …, M}, and

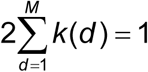

The values of M and σ were explored differently between two network scenarios, fully connected (dense) and sparsely connected networks. In the fully connected network, all neurons were synaptically interconnected, and the spillover kernel *k*(*d*) was normalized by the dispersion of the Hopfield field (the net input) at each neuron, to maintain effect consistency across individual kernels. In memory retrieval simulations, σ values were tested in the range from 0 to 10 while M was fixed at 100 neighbors. The latter reflected the number of neighboring synapses withing a 2-3 µm radius sphere in cortical L1 neuropil with a numerical density of 0.79 µm^−3^ ^55,56^.

In the sparse network, only three neighbors were synaptically connected to each neuron. The extent of excitation spillover was controlled by the number of affected neighbours M (including neurons that are not synaptically connected; illustration in Figure 4d) under a fixed value of its virtual spatial decay, σ = 5.

#### Computing environment

Monte Carlo simulations were run using three computing environments. Firstly, a dedicated 8-node BEOWULF-style diskless PC cluster running under the Gentoo LINUX operating system (kernel 4.12.12), an upgraded version of that described earlier ^99^. Individual nodes comprised an HP ProLiant DL120 G6 Server containing a quad-core Intel Xeon X3430 processor and 8GB of DDR3 RAM. Nodes were connected through a NetGear Gigabit Ethernet switch to a master computer that distributes programs and collects the results on its hard disk. Secondly, on a UCL Myriad cluster: processors for each node, Intel(R) Xeon(R) Gold 6240 CPU @ 2.60GHz; cores per node 36 + 4 A100 GPUs; RAM per node 192GB, tmpfs 1500G, total 6 nodes. Thirdly, cloud computing with Amazon AWS: t4g.medium, memory 4GB. Parallelization and optimization of the algorithms and program codes were implemented by AMC Bridge LLC (Waltham, MA). Hopfield network simulations were carried out using MATLAB R2025b on a standard Windows 11 workstation equipped with a 12th Gen Intel(R) Core(TM) i9-12900K CPU (3.20 GHz, 16 physical cores, 12 logical processors).

## SUPPLEMENTARY FIGURES

**Supplementary figure S1.**
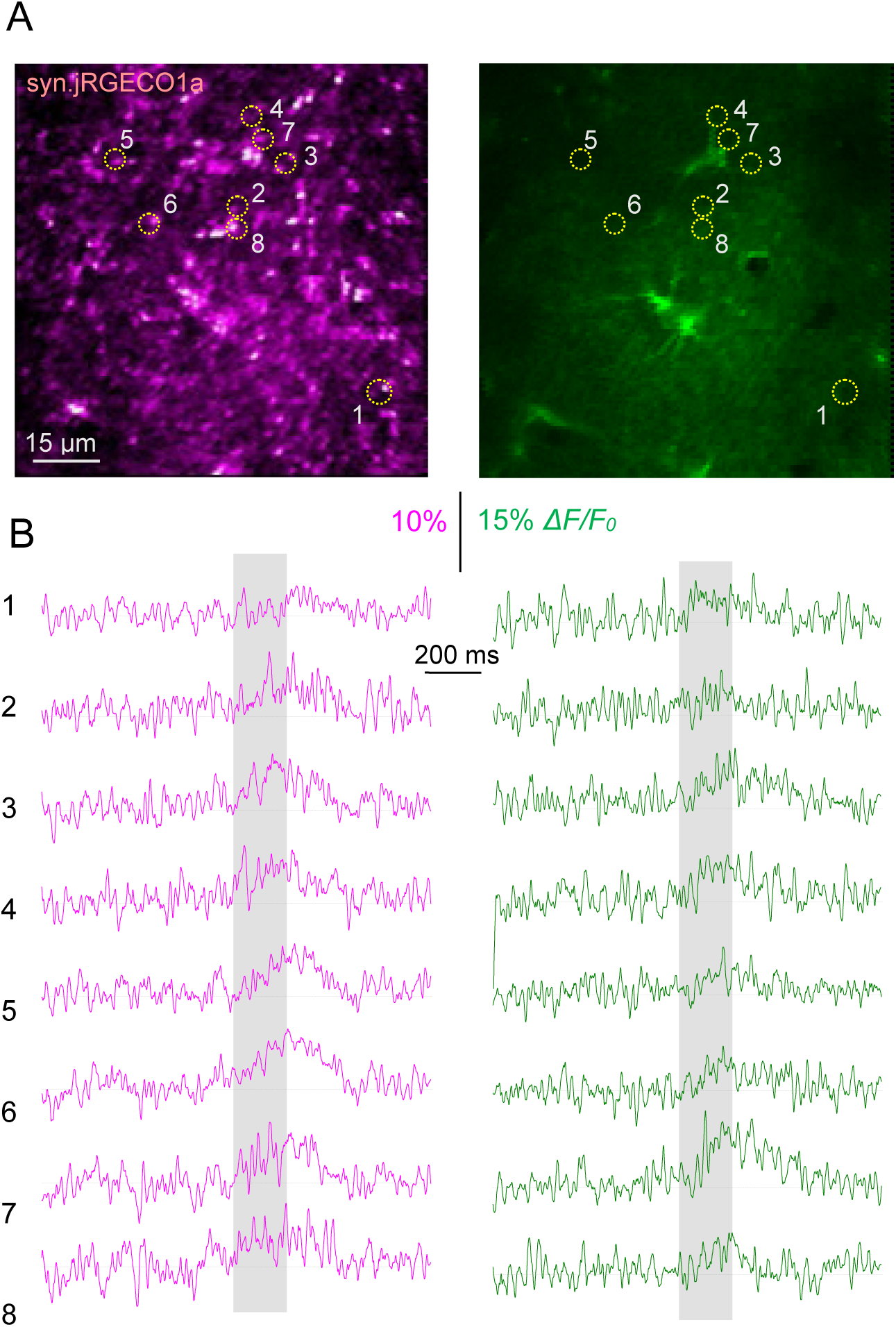
Multiplexed imaging of RWS-evoked responses in hSyn.jGECO1a-expressing thalamocortical axons and GFAP.SF-iGluSnFR expressing astroglia in the barrel cortex. (A) Snapshot example of multiplexed imaging in the jRGECO1a (left) and iGluSnFR (right) channels, with tentative POm axonal boutons (dotted circular ROIs, 1 to 8) selected for analyses based on the detectable presynaptic Ca^2+^ and glutamate release activity. (B) Fluorescence responses in POm axonal boutons shown in A (area-integrated in ROIs 1-8 shown in A) to test RWS (5 s at 3 Hz, grey area), as indicated; frame scan at 30 Hz; single trials (raw data) shown recorded simultaneously in jRGECO1a (left) and iGluSnFR (right) channels.

**Supplementary figure S2.**
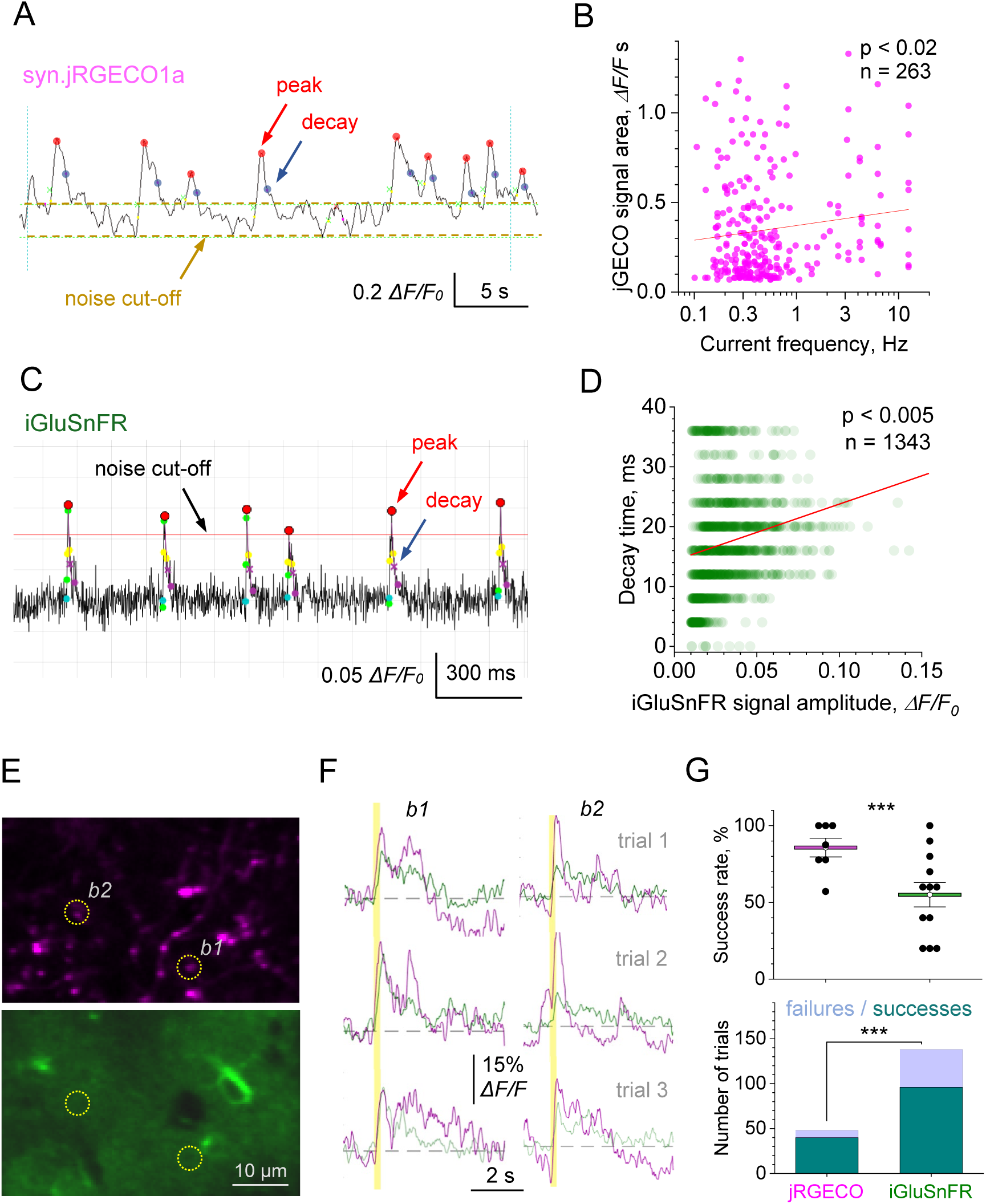
Spontaneous and evoked presynaptic Ca^2+^ activity and glutamate release at POm axonal boutons in the barrel cortex. (A) Example, analysis of spontaneous presynaptic Ca^2+^ activity events (jRGECO1a channel; Mini Analysis 6, Synaptosoft). Main filtering parameters: baseline average window, 100 ms; peak search, 200 ms; decay time search, 500 ms; decay time point, *e*^-1^; detection cut-off, ∼2SD of baseline noise. (B) Magnitude of spontaneous Ca^2+^ activity events (area under the *ΔF/F*_0_ jRGECO1a curve) plotted against the ongoing event frequency (inverse timespan between the current and previous event, n = 263 intervals, 6 boutons); line, linear regression (test, linear-fit ANOVA). (C) Example, analysis of spontaneous synaptic glutamate release events (iGluSnFR channel; Easy Electrophysiology v2.7.3). Main filtering parameters: baseline average window, 30 ms; peak search, 30 ms; decay time search, 150 ms; decay time point, *e*^−1^; detection cut-off, ∼2SD of baseline noise. (D) Decay time of spontaneous glutamate release events (iGluSnFR signal) plotted against signal amplitude (n = 1342 events, 5 boutons); line, linear regression (test, linear-fit ANOVA). (E) Multiplexed imaging of Ca^2+^ activity (jRGECO1a-expressing axonal boutons, top) and glutamate release (iGluSnFR-expressing astroglia channel, bottom) in the barrel cortex; b1 and b2, examples of two axonal boutons showing responses to RWS. (F) Examples of fluorescence responses (magenta jRGECO1a, green iGluSnFR) recorded using resonant scanner frame imaging from boutons b1 and b2 in E, in response to 200 ms RWS at 20 Hz (vertical segment). (G) Top: success rate (proportion of fluorescence responses detected above 3SD noise in all trials), in jRGECO1a (mean ± SEM: 85.7 ± 6.1% success rate, n = 7 boutons, 50 trials) and iGluSnFR channels (55.0 ± 7.9 % success rate, n = 12 boutons, 136 trials); dots, individual experiments (8-12 trials each); ***, p < 0.005 (*F*-test for variance). Bottom: proportions of failures / successes in the two channels, as indicated; ***, p = 0.00678 (Proportion test, Z=2.707).

**Supplementary figure S3.**
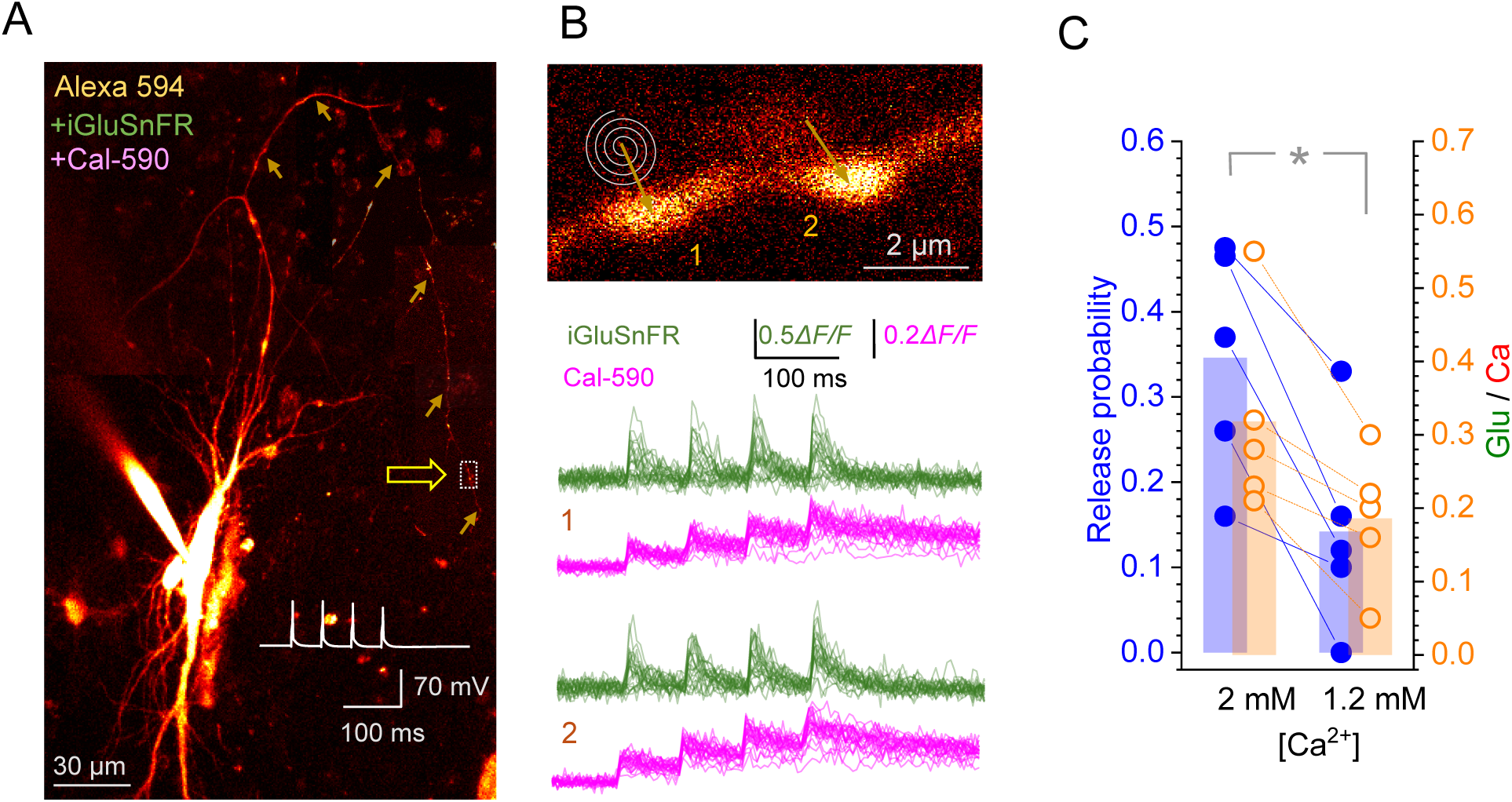
The ratio of evoked glutamate versus Ca^2+^ fluorescence signals at presynaptic boutons tracks changes in release probability: proof-of-principle test. (A) CA3 pyramidal cell (organotypic hippocampal slice) expressing SF-iGluSnFR.A184V and dialysed whole-cell with Cal-590 and 4 μM Alexa-594, with 4 ROIs along the axon (arrowheads, dotted rectangles; collage of 8-17 μm deep image stack projections, λ_x_^2P^ = 800 nm for tracing; inset trace, four action potentials 50 ms apart elicited at the soma in current-clamp; arrow, axonal ROI; see ref. ^1^ for further method detail. (B) Image: Presynaptic axonal boutons (ROI shown in A); spirals and arrows, Tornado scan positions. Traces: Fluorescence time course (integrated Tornado scan signal, 500Hz sampling) in iGluSnFR (green) and Cal-590 (magenta) channels (λ_x_^2P^=910 nm), in response to four action potentials 50 ms apart, as indicated, for the two boutons as shown; 20-24 trials overlaid. Clear distinction between successes and failures at first response provides direct readout of release probability P_r_ as the rate of success. (C) Calculated release probability (green) and Glu/Ca (*ΔF/F_0_* iGluSnFR / *ΔF/F_0_* Cal-590) ratios (orange) at two concentrations of extracellular Ca^2+^, as indicated; dots, individual experiments; bars, average values; *, p < 0.01 (n = 5), in both cases.

**Supplementary figure S4.**
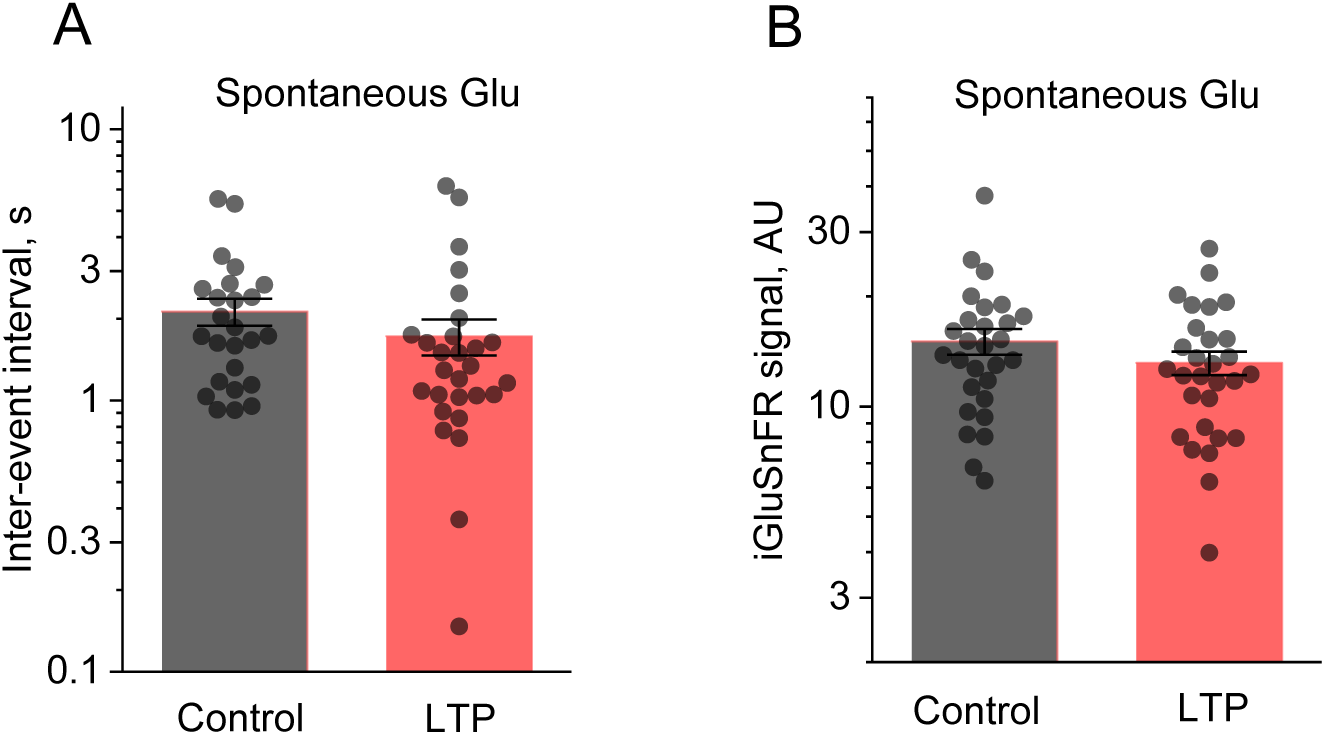
Baseline and spontaneous presynaptic Ca^2+^ (jRGECO1a) and glutamate (iGluSnFR) signals during the induction of RWS LTP. (A) Intervals between successive spontaneous glutamate release events in control conditions and after RWS-LTP induction, as indicated; dots, individual intervals; bars, mean ± SEM (n = 28 and 25 boutons in three animals, respectively; three boutons were lost after LTP induction). (B) Intervals between successive spontaneous glutamate release events in control conditions and after RWS-LTP induction, as indicated; dots, individual intervals; bars, mean ± SEM (n = 29 and 28 boutons in 3 animals, respectively).

**Supplementary figure S5.**
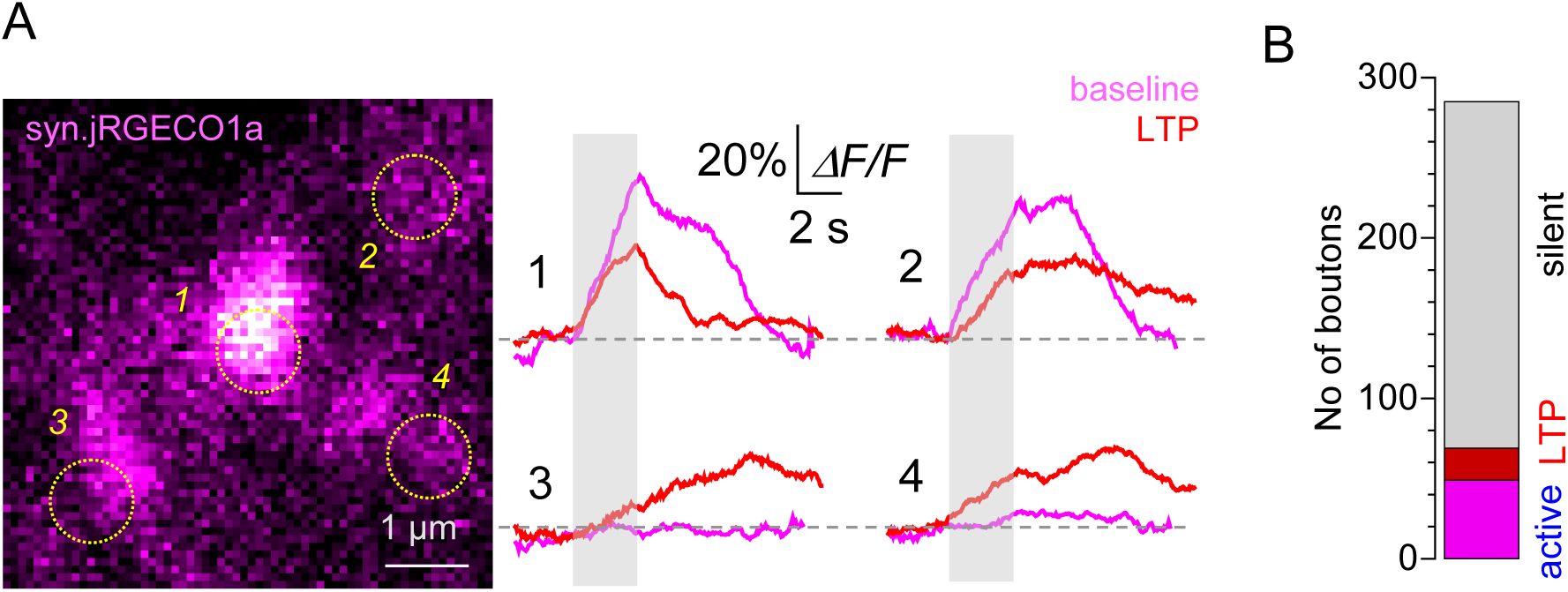
RWS-LTP induction engages previously silent thalamocortical connections in the barrel cortex. (A) Image, example of four arbitrarily selected thalamocortical axonal boutons (jRGECO channel); Traces, boutons 1 and 2 show robust Ca^2+^ response to test RWS (grey segment), which is reduced 10-20 min after RWS-LTP; boutons 3 and 4 show detectable Ca^2+^ response only after RWS-LTP induction. (B) The numbers of thalamocortical axonal boutons that were silent (grey) or RWS-responsive (magenta) throughout the experiment, and RWS-responsive only after LTP induction (red), in individual tested animals (n = 13), as further illustrated in Figure 3E-G.

**Supplementary figure S6.**
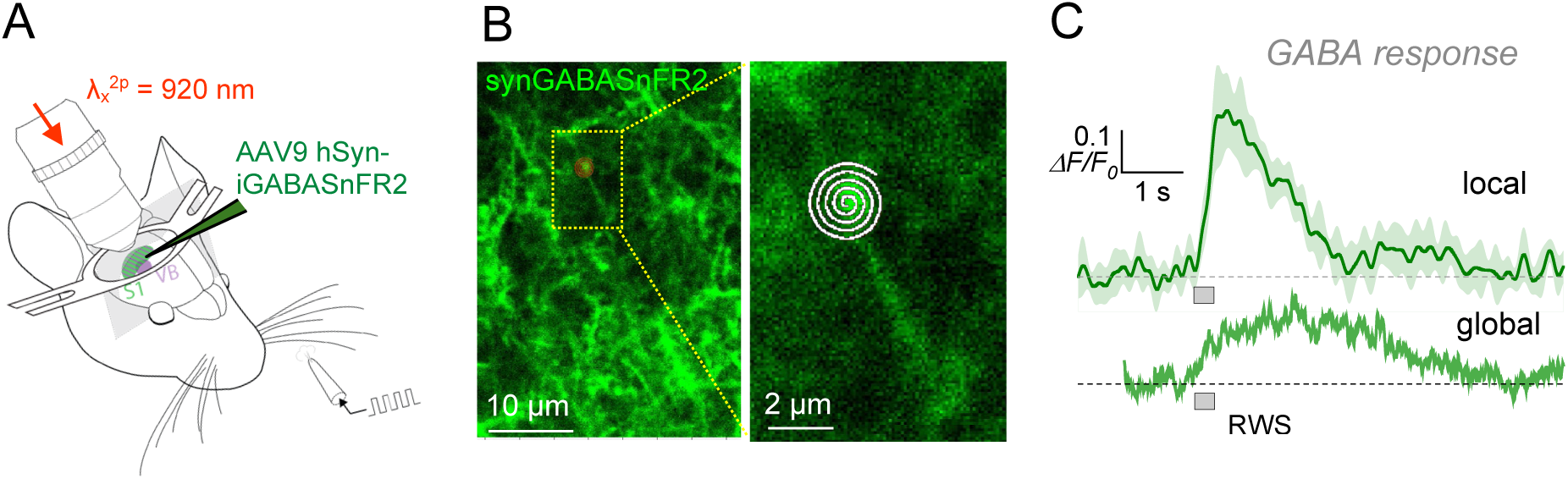
RWS-induced extracellular GABA dynamics monitored with the GABA sensor iGABASnFR2 expressed in cortical neurons. (A) Experimental arrangement: viral expression of syn.iGABASnFR2 in the barrel cortex under the RWS paradigm. (B) Characteristic expression pattern of syn.iGABASnFR2 in neuronal processes in SB1 area (∼110 µm depth, left), with a magnified fragment showing a Tornado scan focusing on a syn.iGABASnFR2-expressing cell process (right). (C) Typical iGABASnFR2 responses to RWS (200 ms at 40 Hz, grey segment) recorded by a Tornado scan (500 Hz) at a local spot shown in B (top trace, mean ± SEM, 3 trials) and over a 120 µm wide local area (bottom trace, single trial).

**Supplementary figure S7.**
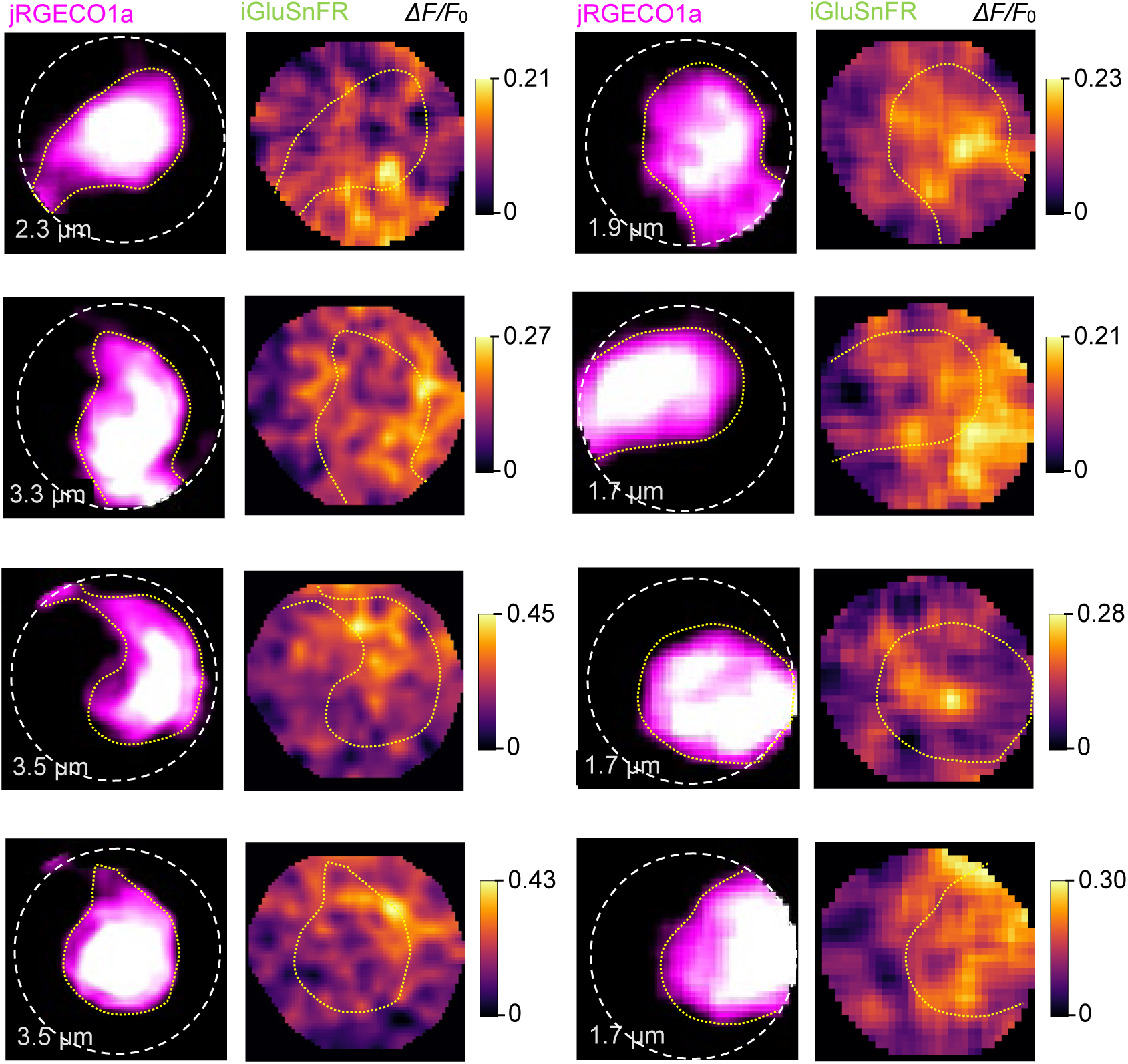
Examples of recorded extrasynaptic glutamate escape in thalamocortical synapses activated by whisker stimulation. Left columns: 2D-reconstructed high-resolution Tornado scans of axonal bouton areas, showing jRGECO1a-expressing boutons; dotted circles, Tornado scan area coverage (diameter shown); yellow dotted line, axonal bouton as judged by the fluorescence of cytosolic jRGECO1a. Right columns (false *ΔF/F_0_* colour scale shown): the *ΔF/F_0_*iGluSnFR signal landscapes, generated as the *ΔF/F_0_* image ratio where *ΔF* and *F*_0_ are the landscape images of astroglia-expressed iGluSnFR signals (green channel) recorded during 200 ms before and 200 ms during a 200 ms long RWS, as illustrated in Fig. 4A-C.

**Supplementary figure S8.**
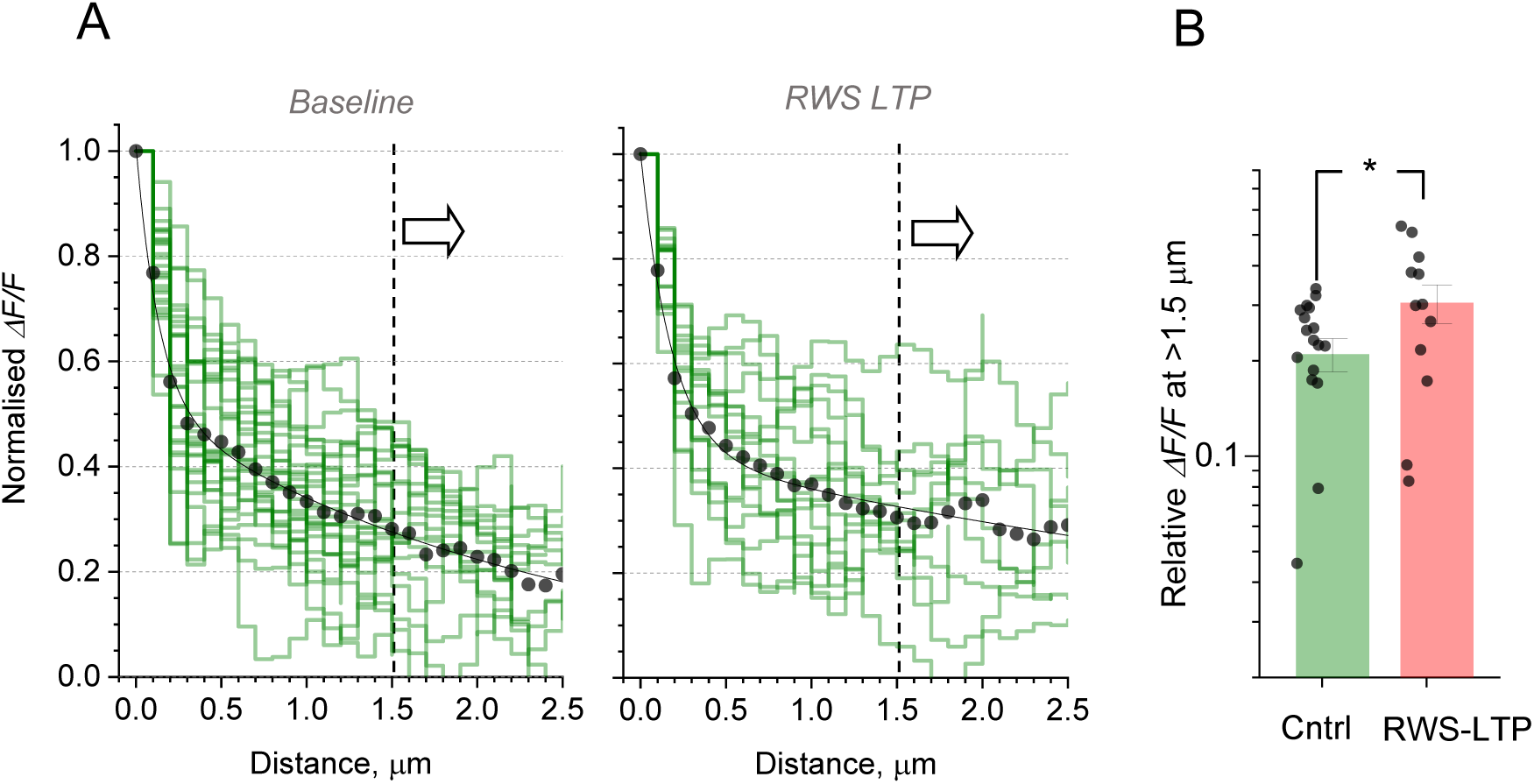
RWS-LTP leads to an increase in glutamate presence at longer distances (> 1.5 µm) from active thalamocortical synapses. (A) Normalised (otherwise unprocessed raw data, green lines) overlaid profiles of the spatial decay displayed by the *ΔF/F_0_* iGluSnFR signal with respect to its hotspot, in baseline conditions (left) and 10-30 min after RWS-LTP induction (right); individual green lines represent individual axonal boutons; dots, average data; solid line, best-fit biexponential approximation, as in Figure 4D; dotted line and arrow indicate the distance range of interest. (B) The *ΔF/F_0_*iGluSnFR relative signal decay averaged over 1.5-2.5 µm distance range (as shown in A) for individual synaptic boutons (dots); columns, mean ± SEM at 0.21 ± 0.03 (n = 18) and 0.31 ± 0.04 (n = 12) for baseline and LTP, respectively; *, p < 0.05, two-sample *t*-test; this analysis ignores boutons in which iGluSnFR signals were not recorded beyond 1.5 µm.

**Supplementary figure S9.**
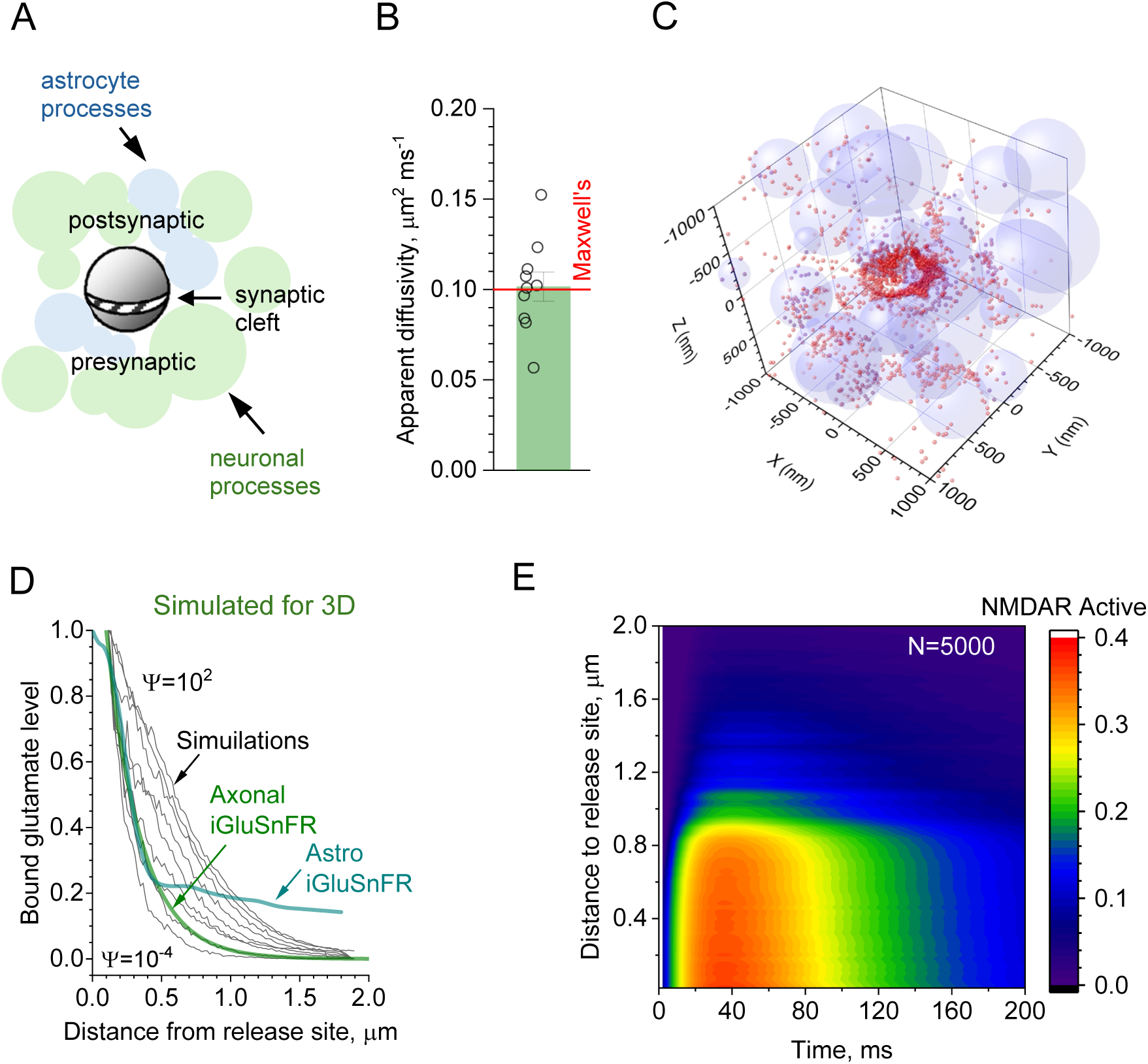
Extrasynaptic glutamate escape simulated in thalamocortical (TC) synapses activated by whisker stimulation. (A) Schematic illustration of the simulated synaptic environment: glutamate molecules are ’released’ instantaneously inside a 220 nm-wide, 20 nm-high synaptic cleft ^2^ which separates two hemispheres representing constant pre-and postsynaptic obstacles to diffusion ^3,4^. The synapse is surrounded by stochastically generated structures that represent either neuronal processes or glutamate transporter-enriched astrocyte processes that occupy, respectively, ∼70% and ∼10% tissue volume fraction in barrel cortex L1 ^5,6^, leaving ∼20% for the extracellular space ^7,8^. These processes are modelled as randomly sized overlapping and concatenated spheres (often forming caterpillar-like structures), as described earlier ^9,10^. Outside the cleft, glutamate diffuses in the tortuous extracellular space with diffusivity *D* = 0.5 μm^2^/ms measured previously with anisotropy-FLIM ^11^. (B) Control tests for the model of glutamate release and free diffusion in a porous neuropil represented by a random scatter of unequal overlapping spheres: comparison of Monte Carlo simulation outcomes (dots, individual trials; bar, mean ± SEM) with the theoretical apparent diffusivity *D_app_* in a porous medium (horizontal line; Maxwell’s equation 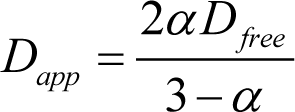 where *α* is medium porosity). Main parameters: simulation arena, 4 μm wide cube; release site, centre; number of Brownian particles 2000; free-space diffusivity *D* = 0.5 μm^2^ /ms; space fraction occupied by spheres, 80% (*α* = 0.2). (C) Snapshot of Monte Carlo simulations (2 x 2 x 2 µm^3^ fragment of the 4 x 4 x 4 µm^3^ arena) showing glutamate molecules (red dots) 4 ms after release inside the synaptic cleft, with a stochastically generated perisynaptic neuropil represented by overlapped concatenated spheroids (as in Figure 4H): for clarity, only astroglial elements are shown. Model parameters represent layer 1 cortical neuropil ^5,6^: ∼20% volume fraction for extracellular space, ∼10% astroglia, ∼70% neuronal elements; see Methods for further detail ^10^. At 4 ms post-release, >90% glutamate molecules are bound to transporter-enriched astrocyte surfaces. (D) Black lines, simulated concentration profiles of glutamate bound to astrocyte surfaces, relative to the hotspot site (grey segment, synaptic cleft area), over the range of astrocyte binding capacity parameter Ψ (10^-4^, 10^-3^, 10^-2^, 0.1, 0.3, 0.5, 0.75, 1, 10, 100 ms); lowest and highest Ψ values shown; green and cyan lines, 3D-reconstructed profiles (Abel conversion) of the RWS-evoked *ΔF/F_0_* signal from POm-and astrocyte -expressed iGluSnFR, respectively, as in Figure 4D and 4F, as indicated. As expected, the signal from astro-iGluSnFR showed a markedly elevated signal beyond the approximate axonal boundary at ∼0.5 µm. (E) Example, dynamics of NMDA receptor double binding by glutamate (to be activated upon membrane depolarization) in simulation experiments illustrated in Figure 4I: 5000 molecules were released instantaneously in the centre of the synaptic cleft and the volume fractions were, in accordance with L1 cortical anatomy ^12^: extracellular space 20%, neuronal elements 70%, astroglia 10%; glutamate binding sites (such as GLT-1 transporters) are expressed throughout astroglial surfaces at a density which corresponds to Ψ = 0.1 ms.

**Supplementary Figure S10.**
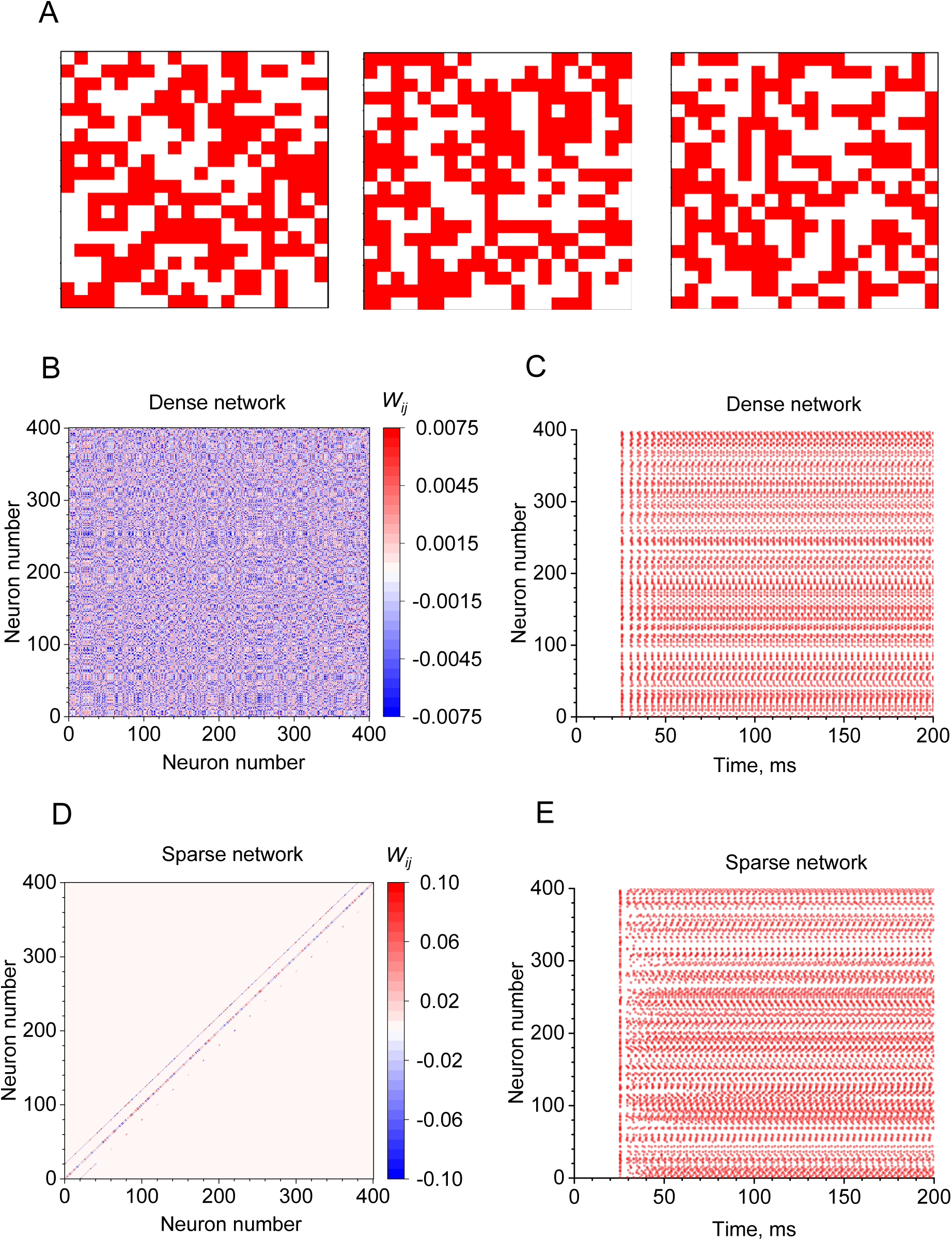
Operation of the fully connected and sparsely connected networks Hopfield’s spiking networks: examples. (A) Examples of three binary patterns that the network was trained to store simultaneously; the network has N = 400 neurons shown as a 20 × 20 grid, with red and white pixels depicting active and inactive neurons, respectively. The networks are expected to retrieve one of these patterns upon presentation of the corresponding noisy probe (cue), as illustrated in Figure 5A. (B) Example of synaptic weight matrix *W* in a dense network (fully connected, as in Figure 5B) upon memorising the patterns shown in A using the Hebbian learning rule. Weights *W_ij_*are dimensionless, normalized Hopfield 1/N units (STAR Methods) representing a connection strength between presynaptic neuron *j* and postsynaptic neuron *i*, varying from negative (blue) to positive (red). In a dense network, almost all elements are non-zero, so the matrix is filled. (C) Cell firing raster plot during memory recall in a dense network displaying spiking activity (each dot represents one action potential) of all 400 neurons while the network retrieves a stored pattern. The memory cue (probe) is applied at time zero, with the recurrent connections driving the network into the stored pattern, producing stable, repeating activity. (D) Example of synaptic weight matrix *W* in a sparse network (Figure 5F) upon memorising the patterns shown in A. As each neuron connects only to its three neighbours, only a few diagonal lines appear, reflecting the grid geometry: neighbours along one direction get neighbouring index numbers (central line), and neighbours along the other direction are 20 numbers apart (outer lines). Other notations as in B. (E) Cell firing raster plot during memory recall in a sparse network. Other notations as in C.

